# Sound improves neuronal encoding of visual stimuli in mouse primary visual cortex

**DOI:** 10.1101/2021.08.03.454738

**Authors:** Aaron M. Williams, Christopher F. Angeloni, Maria N. Geffen

**Affiliations:** Department of Otorhinolaryngology, University of Pennsylvania, Philadelphia, United States; Department of Neuroscience, University of Pennsylvania, Philadelphia, United States; Department of Neurology, University of Pennsylvania, Philadelphia, United States; Department of Psychology, University of Pennsylvania, Philadelphia, United States

**Author notes:** ***Corresponding author*:** Maria Neimark Geffen, Stemmler Hall G10, 3450 Hamilton Walk, Philadelphia, PA 19104.

**Keywords:** audiovisual integration, primary visual cortex, stimulus decoding, electrophysiology

## Abstract

In everyday life, we integrate visual and auditory information in routine tasks such as navigation and communication. Whereas concurrent sound can improve visual perception, the neuronal correlates of this audiovisual integration are not fully understood. Specifically, it remains unclear whether neuronal firing patters in the primary visual cortex (V1) of awake animals demonstrate similar sound-induced improvement in visual discriminability. Furthermore, presentation of sound is associated with movement in the subjects, but little is understood about whether and how sound-associated movement affects audiovisual integration in V1. We investigated how sound and movement interact to modulate V1 visual responses in awake, head-fixed mice and whether this interaction improves neuronal encoding of the visual stimulus. We presented visual drifting gratings with and without simultaneous auditory white noise to awake male and female mice while recording mouse movement and V1 neuronal activity. Sound modulated light-evoked activity of 80% of light-responsive neurons, with 95% of neurons increasing activity when the auditory stimulus was present. Sound consistently enhanced movement, however, a generalized linear model revealed that sound and movement had distinct and complementary effects of the neuronal visual responses. Furthermore, decoding of the visual stimulus from the neuronal activity was improved with sound, even when controlling for movement. Thus, sound and movement modulate visual responses in complementary ways, improving neuronal representation of the visual stimulus. This study clarifies the role of movement as a potential confound in neuronal audiovisual responses and expands our knowledge of how multimodal processing is mediated in the awake brain.

**Significance statement:** Sound and movement are both known to modulate visual responses in the primary visual cortex, however sound-induced movement has largely remained unaccounted for as a potential confound in audiovisual studies in awake animals. Here, authors found that sound and movement both modulate visual responses in an important visual brain area, the primary visual cortex, in distinct, yet complementary ways. Furthermore, sound improved encoding of the visual stimulus even when accounting for movement. This study reconciles contrasting theories on the mechanism underlying audiovisual integration and asserts the primary visual cortex as a key brain region participating in tripartite sensory interactions.

## Introduction

Our brains use incoming sensory information to generate a continuous perceptual experience across sensory modalities. The neuronal systems underlying sensory perception of different modalities interact in a way that often improves perception of the complementary modality (Gingras et al., 2009; Gleiss and Kayser, 2012; Bigelow and Poremba, 2016; Hammond-Kenny et al., 2017; Meijer et al., 2018; Stein et al., 2020). In the audiovisual realm, it is often easiest to understand what someone is saying in a crowded room by additionally relying on visual cues such as lip movement and facial expression (Maddox et al., 2015; Tye-Murray et al., 2016). The McGurk effect and flash-beep illusion are other common perceptual phenomena that demonstrate mutual interactions between the auditory and visual systems (McGurk and MacDonald, 1976; Shams et al. 2002).

The benefits of additional sensory modalities on unisensory processing do not just apply to complex vocal and auditory behavioral interactions. Concurrent sounds such as auditory white noise and pure tones improve sensitivity to and discriminability of visual contrast gradients in humans (Lippert et al., 2007; Chen et al., 2011; Tivadar et al., 2020). The use of these basic audiovisual stimuli has demonstrated that the most robust multisensory perceptual improvements occur around threshold discrimination levels of the otherwise unisensory modality (Chen et al., 2011; Gleiss and Kayser, 2012; Breman et al., 2017). The relative timing of the sensory components is also a factor in their integration. Simultaneous onset and offset of the auditory and visual components strengthens multisensory perceptual improvements compared to asynchronous stimuli (Lippert et al., 2007). Additionally, multisensory integration is often optimal when modulations in visual intensity and auditory amplitude are temporally congruent (Atilgan et al., 2018), likely mimicking covariance of multisensory signals from natural objects. Despite this current understanding of audiovisual integration at a perceptual level, a detailed understanding of the neuronal code that mediates this improvement is still unfolding.

Previous studies of neuronal correlates of audiovisual integration found that the primary sensory cortical areas particupate in this process (Wang et a., 2008; Iurilli et al., 2012; Ibrahim et al., 2016; Meijer et al., 2019; Deneux et al., 2019; McClure and Polack, 2019). The primary visual cortex (V1) contains neurons whose light-evoked firing rates are modulated by sound, as well as neurons that are responsive to sound alone. (Knöpfel et al., 2019). Orientation and directional tuning of individual neurons are also affected by sound. In anesthetized mice, layer 2/3 neurons in V1 exhibited sharpened tuning in the presence of sound (Ibrahim et al., 2016), providing a potential mechanism through which sound improves visual encoding. Whereas initial studies found a suppressive signal provided by the primary auditory cortex to V1 (Iurilli et al., 2012; Ibrahim et al., 2016), later studies found heterogeneous changes across neurons in visual tuning curve bandwidth with and without sound (Meijer et al., 2017; McClure and Polack, 2019). These contrasting findings raise the question of whether previously reported sound-induced changes in V1 neuronal activity in awake animals results in improved visual processing, and through which coding schemes these effects are mediated. Ultimately, this hypothesized improvement in visual encoding would provide a missing link between cross-sensory neuronal responses and the field’s current understanding of behavioral and perceptual effects described above.

An important factor that has thus far been largely unaccounted for in audiovisual studies is that awake animals are subject to brain-wide changes in neuronal activity due to stimulus-aligned, uninstructed movements (Musall et al., 2019). Sound-induced movement represents a potential confound for audiovisual studies in awake animals because whisking and locomotion modulate neuronal activity in the sensory cortical areas. In V1, movement enhances neuronal visual responses and improves neuronal encoding of the visual scene (Niell and Stryker, 2010; Dadarlat and Stryker, 2017). Conversely, in the auditory cortex (AC), locomotion suppresses neuronal spontaneous and auditory responses (Nelson et al., 2013; Schneider and Mooney, 2018; Bigelow et al., 2019). Therefore, the contribution of movement to neuronal responses to multi-sensory stimuli is likely due to multiple processes and can greatly affect audiovisual integration.

Thus, audiovisual integration in V1 may not simply represent afferent information from auditory brain regions. Whereas V1 neurons are sensitive to the optogenetic stimulation (Ibrahim et al., 2016) and pharmacologic suppression (Deneux et al., 2019) of AC neurons, the modulation of V1 activity may instead be a byproduct of uninstructed sound-induced movements which themselves modulate visual responses (Bimbard et al., 2021). Here, we tested these alternative explanations of the extent to which movement contributes to audiovisual integration in V1 by performing extracellular recordings of neuronal activity in V1 while monitoring movement in awake mice presented with audiovisual stimuli. We used these results to build on prior studies reporting sound-induced changes in V1 visual responses (Ibrahim et al., 2016; Meijer et al., 2017; McClure and Polack, 2019), in order to determine whether and through what coding mechanism this cross-modal interaction improves visual encoding in awake mouse subjects. The audiovisual stimulus consisted of auditory white noise and visual drifting gratings in order to allow comparison of sound’s effect across the visual contrast parameter. We found that the majority of neurons in V1 were responsive to visual and auditory stimuli. Sound and movement exerted distinct yet complementary effects on shaping the visual responses. Importantly, sound improved discriminability of the visual stimuli both in individual neurons and at a population level, an effect that persisted when accounting for movement.

## Materials and methods

### Mice

All experimental procedures were in accordance with NIH guidelines and approved by the IACUC at the University of Pennsylvania. Mice were acquired from Jackson Laboratories (5 male, 6 female, aged 10-18 weeks at time of recording; B6.Cast-*Cdh23*^*Ahl+*^ mice [Stock No: 018399]) and were housed at 28°C in a room with a reversed light cycle and food provided ad libitum. Experiments were carried out during the dark period. Mice were housed individually after headplate implantation. Euthanasia was performed using CO_2_, consistent with the recommendations of the American Veterinary Medical Association (AVMA) Guidelines on Euthanasia. All procedures were approved by the University of Pennsylvania IACUC and followed the AALAC Guide on Animal Research. We made every attempt to minimize the number of animals used and to reduce pain or discomfort.

### Surgical procedures

Mice were implanted with skull-attached headplates to allow head stabilization during recording, and skull-penetrating ground pins for electrical grounding during recording. The mice were anesthetized with 2.5% isoflurane. A ∼1mm craniotomy was performed over the right frontal cortex, where we inserted a ground pin. A custom-made stainless steel headplate (eMachine Shop) was then placed on the skull at midline, and both the ground pin and headplate were fixed in place using C&B Metabond dental cement (Parkell). Mice were allowed to recover for 3 days post-surgery before any additional procedures took place.

### Electrophysiological recordings

All recordings were carried out inside a custom-built acoustic isolation booth. 1-2 weeks following the headplate and ground pin attachment surgery, we habituated the mice to the recording booth for increasing durations (5, 15, 30 minutes) over the course of 3 days. On the day of recording, mice were placed in the recording booth and anesthetized with 2.5% isoflurane. We then performed a small craniotomy above the left primary visual cortex (V1, 2.5mm lateral of midline, 0-0.5 mm posterior of the lambdoid suture). Mice were then allowed adequate time to recover from anesthesia. Activity of neurons were recorded using a 32-channel silicon probe (NeuroNexus A1×32-Poly2-5mm-50s-177). The electrode was lowered into the primary visual cortex via a stereotactic instrument to a depth of 775-1000μm. Following the audiovisual stimulus presentation, electrophysiological data from all 32 channels were filtered between 600 and 6000 Hz, and spikes belonging to single neurons and multi-units were identified in a semi-automated manner using KiloSort2 (Pachitariu et al., 2016).

### Audiovisual stimuli

The audiovisual stimuli were generated using MATLAB (MathWorks, USA), and presented to mice on a 12” LCD monitor (Eyoyo) with a 60Hz framerate and through a magnetic speaker (Tucker-Davis Technologies) placed to the right of the mouse. The visual stimulus was generated using the PsychToolBox package for MATLAB and consisted of square wave drifting gratings 1 s in duration, 4-Hz temporal frequency, and 0.1 cycles/°. The gratings moved in 12 directions, evenly spaced 0°-360°, and were scaled to a range of 5 different visual contrast levels (0, 0.25, 0.5, 0.75, 1), totaling 60 unique visual stimuli. The auditory stimulus was sampled at 400 kHz and consisted of a 1 s burst of 70 dB white noise. The visual grating was accompanied by the auditory noise on half of trials (120 unique trial types, 10 repeats each), with simultaneous onset and offset. A MATLAB-generated TTL pulse aligned the onset of the auditory and visual stimuli, and was verified using a ThorLabs photodetector and microphone. This TTL pulse was also used to align the electrophysiological recording data with the audiovisual stimulus trials. The auditory-only condition corresponded to the trials with a visual contrast of 0. The trial order was randomized and was different for each recording.

### Data analysis and statistical procedures

Spiking data from each recorded unit was organized by trial type and aligned to the trial onset. The number of spikes during each trial’s first 0-300ms was input into a generalized linear model (GLM; predictor variables: visual contrast [continuous variable 0, 0.25, 0.5, 0.75, 1], sound [0 or 1]; response variable: number of spikes during 0-300ms; Poisson distribution, log link function), allowing the classification of each neuron’s responses as having a main effect (p<0.05) of light, sound, and/or a light-sound interaction. Neurons that were responsive to both light and sound or had a significant light-sound interaction term were classified as “light-responsive sound-modulated.” To quantify the supra- or sub-linear integration of the auditory and visual responses, we calculated the linearity ratio of neurons’ audiovisual responses. This ratio was defined as FR_AV_ / (FR_V_ + FR_A_), and the sound-only response FR_A_ was calculated using the trials with a visual contrast of 0.

We calculated mutual information (MI) between neuronal responses and the five different visual contrast level, as well as between neuronal responses and the 12 different drifting grating directions, in order to guide the response time window used for our subsequent analyses. We calculated mutual information according to the equations (Borst and Theunissen, 1999):

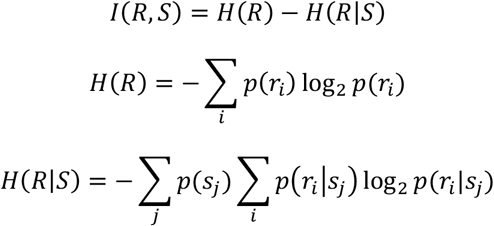

where I(R,S) is the MI between the neuronal response R and visual stimulus S, H(R) is the entropy of neuronal response R, and H(R|S) is the entropy of neuronal response R given the stimulus S. S_j_ represents the stimulus parameter either visual contrast or grating direction, and r_i_ represents the number of spikes in a specific time window. We used a sliding 10ms time window to serially calculate MI with the visual stimulus across the neuronal response. We then averaged the MI trace across neurons to generate a population mean trace.

We quantified changes in response timing by calculating response latency, onset slope, and onset response duration. First, mean peristimulus time histograms (PSTH) were constructed for each trial type using a 10 ms sliding window. The latency was calculated as the first time bin after stimulus onset in which the mean firing rate at full contrast exceeded 1 standard deviation above baseline. The slope Hz/ms slope was calculated from the trial onset to the time of the peak absolute value firing rate. The response duration was calculated using the full width at half maximum of the peak firing rate at stimulus onset (limited to 0-300 ms).

Orientation selectivity and direction selectivity were determined for all light-responsive neurons. The preferred direction of each direction-selective neuron was found by calculating half the complex phase (Neill and Stryker, 2008) of the value

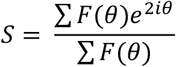

We calculated orientation and direction-selective indices (Zhao et al., 2013) for each neuron according to:

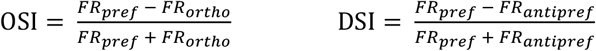

where FR_ortho_ and FR_antipref_ are the mean firing rates in the orthogonal (90°) and anti-preferred (180°) directions, respectively. One-tailed permutation testing was performed by comparing these OSI and DSI values to pseudo OSI and DSI values obtained by 200 random shuffles of the firing rates from the pooled preferred and orthogonal or anti-preferred trials. If a neuron’s actual OSI or DSI value was >75% of shuffled OSI or DSI values, the neuron was classified as “orientation-” or “direction-selective,” respectively. To determine whether there were statistically significant changes in the preferred direction from the visual to audiovisual conditions, we applied a bootstrapping procedure, subsampling the visual trials for each neuron 1000 times and creating a confidence interval of the mean shift in preferred direction (degrees) for each population randomization.

We assessed and controlled for sound-induced movement as a potential confound for the audiovisual effects observed. During a subset of V1 recordings (9 recordings, 5 mice), mouse movement was tracked throughout stimulus presentation. Video recording was performed using a Raspberry Pi 4 Model B computer system with an 8MP infrared Raspberry Pi NoIR Camera V2 attachment. The camera was positioned to the front and left of the mice, which allowed capture of primarily the forepaw and whisking motion but with more limited hindpaw motion visualization. The video was converted to MP4 format, and motion was then quantified using the freely available Facemap software (Stringer et al., 2019). Within the Facemap GUI, we identified regions of interest (ROIs) on the whiskers and face to capture whisking behavior, ROIs on the extremities to capture locomotive behavior, and ROIs distributed across the mouse subject to capture general, nonspecific movement, which captured both whisking and locomotion. The separate motion energy output from each regions was then aligned to the trials of the audiovisual stimulus from the recording trials for further analysis.

Similar to above, a GLM (predictor variables: visual contrast level, sound presence, average motion during each trial [using the general nonspecific movement trace]; response variable: trial spikes during 0-300ms; Poisson distribution, log link function) classified each neuron as having a main effect (p<0.05) of light, sound, or motion, as well as the pairwise interactions of these parameters. Light-responsive sound-modulated neurons, according to the above definition, that additionally displayed either a main effect of motion or significant light-motion or sound-motion interaction terms were classified as “motion-modulated” and were included for further analysis.

We also visualized the overall distribution of mouse subject movement across trials by calculating a z-score for each trial. Using the nonspecific mouse movement value from each trial, we grouped together trials from each recording session, subtracted the group average, and divided by the group standard deviation in order to obtain a z-score for each trial. This z-score represented whether the mouse moved more or less compared to other trials from that recording session.

In order to reconstruct peristimulus time histograms of light-responsive, sound-modulated, motion-modulated neurons, we used a separate GLM. Using a 10ms sliding window across all trials, we input the visual contrast level, sound presence, and general nonspecific movement during that window (discretized into five bins) as predictor variables, and the number of spikes during that window as response variables, into the GLM (Poisson distribution, log link function) to calculate coefficients for light, sound, motion, and their pairwise interactions. This approach allowed us to reconstruct the mean PSTH of individual neurons observed during each trial type by calculating:

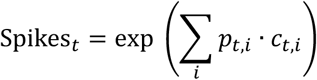

where the spikes in time window *t* are determined by the values *p* and coefficients *c* of predictor variable *i*. From there, we used this same equation to estimate the shape of the PSTHs when varying sound and motion in order to determine differential effects these parameters had on the temporal trajectory of neurons’ visual responses. In a separate analysis, we used a similarly structured GLM, but replaced the “general nonspecific movement” predictor variable with independent locomotion and whisking variables, using the Facemap output from the locomotion- and whisking-related ROIs. This allowed us to additionally report how locomotion and whisking individually modulate visual responses, as opposed to grouped into a single nonspecific movement variable.

The *d’* sensitivity index (Stanislaw and Todorov, 1999; von Trapp et al., 2016) was used to calculate the directional discriminability of direction-selective neurons. The *d’* sensitivity index between two directions θ_1_ and θ_2_ is calculated as:

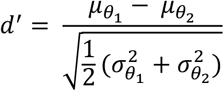

where *μ*_*θ*_ and *σ*_*θ*_ are the response mean and standard deviation, respectively, for direction θ. For each neuron, the sensitivity index was calculated in a pairwise manner for preferred direction versus all other directions and then aligned relative to the preferred direction in order to test sensitivity index as a function of angular distance from preferred direction.

We used a maximum likelihood estimate approach (Montijn et al., 2014; Meijer et al., 2017) to decode the visual stimulus direction from the neuronal responses based on Bayes rule:

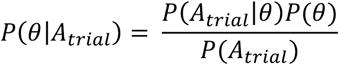

For decoding using individual neurons, the likelihood P(A_trial_|θ) for each orientation or direction was computed based on the Poisson response distribution across all trials of that orientation or direction, with a leave-one-out cross-validation technique in which the probe trial (A_trial_) was excluded from the training data. The prior P(θ) was uniform, and the normalization term P(A_trial_) was similarly applied to all directions. Therefore, the posterior probability P(θ|A_trial_) was proportional to and based on evaluating the likelihood function at the value of the probe trial. For orientation-selective neurons, decoding was performed between the preferred and orthogonal orientations, and for direction-selective neurons, decoding was performed between the preferred and anti-preferred directions. For decoding using populations of neurons, neurons were pooled across recording sessions. A similar approach was used; however, here, the posterior probability P(θ|A_pop_) was proportional to the joint likelihood P(A_pop_|θ) of the single-trial activity across all N neurons in the population (A_pop_):

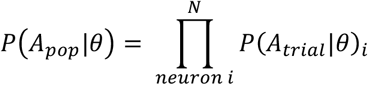

With this population-based analysis, pairwise decoding was performed between every orientation and its orthogonal orientation (1 of 2 options), as well as decoding one direction from all possible directions (1 of 12 options).

Additionally, we used a support vector machine (SVM) to corroborate the findings of the MLE-based decoder. The SVM was implemented using MATLAB’s fitcsvm function with a linear kernel to predict the drifting grating direction based on single-trial population responses. Similarly, a leave-one-out cross-validation technique was used, and pairwise decoding was performed between every combination of two stimulus directions.

### Statistics

Figure data are displayed as means with standard error of the mean (SEM), unless otherwise noted. Shapiro-Wilk tests were used to assess normality, and the statistical tests performed are indicated in the text, figures, and Table 1. For multi-group and multivariate analysis (e.g., ANOVA and Kruskal-Wallis tests) in which a significant (p<0.05) interaction was detected, we subsequently performed a post hoc Bonferroni-corrected test. P-values reported as 0 are too small to be accurately calculated by Matlab (p<2.2e-301), due to characteristically large data sets. See Table 1 for a detailed summary of statistical results and post hoc comparisons.

**Table 1:** Statistical comparisons.

| Comparison | Fig | Test | Test statistic | N | df | p-value | Post hoc test | Post hoc $\alpha$ | Post hoc comparison | Post hoc p-value |
| --- | --- | --- | --- | --- | --- | --- | --- | --- | --- | --- |
| Orientation selective index | 2C | t-test | t-stat = -1.0 | $n_{1000ms}=303$<br>$n_{300ms}=269$ | 565 | p = 0.30 | | | | |
| Direction selective index | 2C | t-test | t-stat = -1.6 | $n_{1000ms}=143$<br>$n_{300ms}=144$ | 281 | p = 0.10 | | | | |
| Mean firing rate, V vs AV | 3C | Paired 2-way ANOVA | F(vis)=340<br>F(aud)=506<br>F(interact)=75 | 565 neurons | vis=4<br>aud=1<br>interact = 4 | p(vis) = 1.2e-100<br>p(aud) = 1.6e-88<br>p(interact) = 5.7e-4 | Bonferroni-corrected paired t-test | 0.01 | Contrast 0, V vs AV | 2.1e-50 |
|  |  |  |  |  |  |  |  |  | Contrast 0.25, V vs AV | 2.6e-62 |
|  |  |  |  |  |  |  |  |  | Contrast 0.5, V vs AV | 5.7e-75 |
|  |  |  |  |  |  |  |  |  | Contrast 0.75, V vs AV | 1.1e-81 |
|  |  |  |  |  |  |  |  |  | Contrast 1, V vs AV | 2.0e-81 |
| Linearity ratio, V vs AV | 3E | Kruskal-Wallis test | Chi-sq = 61 | 555 neurons | 4 | p = 1.6e-12 | Bonferroni-corrected Wilcoxon signed rank test | 0.0125 | Contrast 0 vs 0.25 | 0.053 |
|  |  |  |  |  |  |  |  |  | Contrast 0 vs 0.5 | 0.0040 |
|  |  |  |  |  |  |  |  |  | Contrast 0 vs 0.75 | 4.6e-8 |
|  |  |  |  |  |  |  |  |  | Contrast 0 vs 1 | 2.1e-5 |
| Orientation selectivity index, V vs AV | 4E | Paired t-test | t-stat = 4.8 | 269 neurons | 268 | p = 2.4e-6 |  |  |  |  |
| Direction selectivity index, V vs AV | 4F | Paired t-test | t-stat = 3.5 | 144 neurons | 143 | p = 6.4e-4 |  |  |  |  |
| Onset response latency, V vs AV | 5B | Paired 2-way ANOVA | F(vis)=5.7<br>F(aud)=64<br>F(interact)=2.7 | 517 neurons | vis=3<br>aud=1<br>interact=3 | p(vis)=6.9e-4<br>p(aud)=6.8e-18<br>p(interact)=0.045 | Bonferroni-corrected paired t-test | 0.01 | Contrast 0.25, V vs AV | 2.3e-4 |
|  |  |  |  |  |  |  |  |  | Contrast 0.5, V vs AV | 7.1e-12 |
|  |  |  |  |  |  |  |  |  | Contrast 0.75, V vs AV | 4.6e-5 |
|  |  |  |  |  |  |  |  |  | Contrast 1, V vs AV | 9.9e-4 |
| Onset response slope, V vs AV | 5D | Paired 2-way ANOVA | F(vis)=70<br>F(aud)=66<br>F(interact)=2.8 | 563 neurons | vis=3<br>aud=1<br>interact=3 | p(vis)=3.5e-121<br>p(aud) = 2.7e-15<br>p(interact) = 0.038 | Bonferroni-corrected paired t-test | 0.01 | Contrast 0.25, V vs AV | 1.4e-4 |
|  |  |  |  |  |  |  |  |  | Contrast 0.5, V vs AV | 8.9e-13 |
|  |  |  |  |  |  |  |  |  | Contrast 0.75, V vs AV | 3.6e-12 |
|  |  |  |  |  |  |  |  |  | Contrast 1, V vs AV | 5.5e-8 |
| Onset response duration, V vs AV | 5F | Paired 2-way ANOVA | F(vis)=17<br>F(aud)=129<br>F(interact)=1.4 | 367 neurons | vis=3<br>aud=1<br>Interact=3 | p(vis)=1.3e-10<br>p(aud) = 8.7e-98<br>p(interact) = 0.23 |  |  |  |  |
| Response coefficient of variation, V vs AV | 5H | Paired 2-way ANOVA | F(vis)=1.3<br>F(aud)=834<br>F(interact)=1.0 | 564 neurons | vis=4<br>aud=1<br>Interact=4 | p(vis) = 0.28<br>p(aud) = 4.2e-103<br>p(interact) = 0.38 |  |  |  |  |
| Sound induced movement | 6D | Paired t-test | t-stat = -14.9 | 9 recording sessions | 8 | p = 4.0e-7 |  |  |  |  |
| Firing rate across movement range, V vs AV | 6G | Unbalanced 2-way ANOVA | F(motion)=18<br>F(sound)=32<br>F(interact)=17 | Variable trial count | mot=2<br>aud=1<br>Interact=2 | p(motion) = 1.6e-8<br>p(sound) = 1.6e-8<br>p(interact) = 3.1e-8 | Bonferroni corrected two-sample t-test | 0.016 | Stationary, V vs AV<br>Low motion, V vs AV<br>High motion, V vs AV | 1.0e-15<br>3.1e-10<br>0.59 |
| GLM PSTH, light vs light/sound | 7F | Paired t-test | 1391 unique t-stats | 343 neurons | 342 | 1391 unique p-values,<br>$\alpha = 0.05/1391 = 3.6e-5$ | | | | |
| GLM PSTH, light vs light/motion | 7G | Paired t-test | 1391 unique t-stats | 343 neurons | 342 | 1391 unique p-values,<br>$\alpha = 0.05/1391 = 3.6e-5$ | | | | |
| GLM PSTH, light vs light/sound/motion | 7H | Paired t-test | 1391 unique t-stats | 343 neurons | 342 | 1391 unique p-values,<br>$\alpha = 0.05/1391 = 3.6e-5$ | | | | |
| Orientation decoding accuracy, individual neurons, V vs AV | 8E | Paired 2-way ANOVA | F(vis)=73<br>F(aud)=50<br>F(interact)=7.2 | 264 neurons | vis=4<br>aud=1<br>interact=4 | p(vis)=4.8e-112<br>p(aud)=1.7e-11<br>p(interact) = 1.0e-5 | Bonferroni-corrected paired t-test | 0.01 | Contrast 0, V vs AV<br>Contrast 0.25, V vs AV<br>Contrast 0.5, V vs AV<br>Contrast 0.75, V vs AV<br>Contrast 1, V vs AV | 0.78<br>1.5e-4<br>2.2e-11<br>0.21<br>1.4e-6 |
| Direction decoding accuracy, individual neurons, V vs AV | 8G | Paired 2-way ANOVA | F(vis)=20<br>F(aud)=12<br>F(interact)=0.39 | 140 neurons | vis=4<br>aud=1<br>interact=4 | p(vis)=1.2e-14<br>p(aud)=6.9e-4<br>p(interact)=0.82 |  |  |  |  |
| Orientation decoding accuracy, MLE, population, V vs AV | 9D | 2-way ANOVA | F(vis)=720<br>F(aud)=2.8<br>F(interact)=26 | 50 repeats | vis=4<br>aud=1<br>interact=4 | p(vis) = 2.6e-98<br>p(aud) = 0.098<br>p(interact) = 1.7e-82 | Bonferroni-corrected paired t-test | 0.01 | Contrast 0, V vs AV<br>Contrast 0.25, V vs AV<br>Contrast 0.5, V vs AV<br>Contrast 0.75, V vs AV<br>Contrast 1, V vs AV | 2.1e-7<br>1.5e-12<br>4.1e-9<br>1.4e-4<br>9.4e-4 |
| Direction decoding accuracy, MLE, population, V vs AV | 9E | 2-way ANOVA | F(vis)=99<br>F(aud)=0.03<br>F(interact)=7.8 | 50 repeats | vis=4<br>aud=1<br>interact=4 | p(vis) = 4.2e-90<br>p(aud) = 0.87<br>p(interact) = 8.1e-6 | Bonferroni-corrected paired t-test | 0.01 | Contrast 0, V vs AV<br>Contrast 0.25, V vs AV<br>Contrast 0.5, V vs AV<br>Contrast 0.75, V vs AV<br>Contrast 1, V vs AV | 0.21<br>5.8e-4<br>2.5e-4<br>0.48<br>0.97 |
| Overall decoding accuracy, MLE, population, V vs AV | 9H | 2-way ANOVA | F(vis)=137<br>F(aud)=13<br>F(interact)=5.5 | 20 repeats | vis=4<br>aud=1<br>interact=4 | p(vis)=8.7e-55<br>p(aud)=4.2e-4<br>p(interact)=3.3e-4 | Bonferroni - corrected paired t-test | 0.01 | Contrast 0, V vs AV<br>Contrast 0.25, V vs AV<br>Contrast 0.5, V vs AV<br>Contrast 0.75, V vs AV<br>Contrast 1, V vs AV | 0.011<br>2.9e-4<br>0.090<br>0.0054<br>0.57 |
| Orientation decoding accuracy, individual neurons, V vs AV | 10B | Paired 2-way ANOVA | F(vis) = 80<br>F(aud) = 18<br>F(interact) = 1.2 | 90 neurons | vis=4<br>aud=1 | p(vis) = 1.2e-80<br>p(aud)=5.4e-5<br>p(interact)=0.32 |  |  |  |  |
|  |  |  |  |  | interact=4 |  |  |  |  |  |
| Orientation decoding accuracy, individual neurons, V vs motion-corrected AV | 10B | Paired 2-way ANOVA | F(vis) = 82<br>F(aud) = 8.1<br>F(interact) = 1.7 | 90 neurons | vis=4<br>aud=1<br>interact=4 | p(vis)=6.2e-81<br>p(aud)=5.3e-3<br>p(interact)=0.15 |  |  |  |  |
| Orientation decoding accuracy, individual neurons, V vs sound-corrected AV | 10B | Paired 2-way ANOVA | F(vis) = 70<br>F(aud) = 1.5<br>F(interact) = 1.9 | 90 neurons | vis=4<br>aud=1<br>interact=4 | p(vis) = 7.7e-72<br>p(aud) = 0.23<br>p(interact) = 0.11 |  |  |  |  |
| Population decoding accuracy, V vs AV | 10D | 2-way ANOVA | F(vis) = 337<br>F(aud) = 133<br>F(interact) = 4.2e-10 | 10 repeats | vis=4<br>aud=1<br>interact=4 | p(vis)=3.1e-53<br>p(aud)=2.0e-19<br>p(interact)=4.2e-10 | Bonferroni-corrected paired t-test | 0.01 | Contrast 0, V vs AV | 0.041 |
|  |  |  |  |  |  |  |  |  | Contrast 0.25, V vs AV | 5.6e-6 |
|  |  |  |  |  |  |  |  |  | Contrast 0.5, V vs AV | 3.5e-6 |
|  |  |  |  |  |  |  |  |  | Contrast 0.75, V vs AV | 7.0e-7 |
|  |  |  |  |  |  |  |  |  | Contrast 1, V vs AV | 1.4e-4 |
| Population decoding accuracy, V vs motion-corrected AV | 10D | 2-way ANOVA | F(vis) = 230<br>F(aud) = 74<br>F(interact) = 6.7 | 10 repeats | vis=4<br>aud=1<br>interact=4 | p(vis)=2.5e-46<br>p(aud)=2.1e-13<br>p(interact)=9.3e-5 | Bonferroni-corrected paired t-test | 0.01 | Contrast 0, V vs motion-corrected AV | 0.94 |
|  |  |  |  |  |  |  |  |  | Contrast 0.25, V vs motion-corrected AV | 1.1e-4 |
|  |  |  |  |  |  |  |  |  | Contrast 0.5, V vs motion-corrected AV | 0.01 |
|  |  |  |  |  |  |  |  |  | Contrast 0.75, V vs motion-corrected AV | 0.0023 |
|  |  |  |  |  |  |  |  |  | Contrast 1, V vs motion-corrected AV | 0.021 |
| Population decoding accuracy, V vs sound-corrected AV | 10D | 2-way ANOVA | F(vis) = 192<br>F(aud) = 0.02<br>F(interact) = 9.3 | 10 repeats | vis=4<br>aud=1<br>interact=4 | p(vis)=3.3e-43<br>p(aud) = 0.88<br>p(interact) = 3.4e-6 | Bonferroni-corrected paired t-test | 0.01 | Contrast 0, V vs motion-corrected AV | 0.87 |
|  |  |  |  |  |  |  |  |  | Contrast 0.25, V vs motion-corrected AV | 0.039 |
|  |  |  |  |  |  |  |  |  | Contrast 0.5, V vs motion-corrected AV | 0.19 |
|  |  |  |  |  |  |  |  |  | Contrast 0.75, V vs motion-corrected AV | 0.080 |
|  |  |  |  |  |  |  |  |  | Contrast 1, V vs motion-corrected AV | 0.0025 |

## Results

### Sound enhances the light-evoked firing rate of a subset of V1 neurons

Previous work identified that sound modulates visual responses in V1 (Ibrahim et al., 2016; Meijer et al., 2018; McClure and Polack, 2019), yet how that interaction affects stimulus encoding in individual neurons and as a population in the awake brain is still being revealed. Furthermore, whether that interaction can be exclusively attributed to sound or to sound-induced motion is controversial (Bimbard et al., 2021). To elucidate the principles underlying audiovisual integration, we presented audiovisual stimuli to awake mice while performing extracellular recordings in V1 (Figure 1A). The visual stimulus consisted of drifting gratings in 12 directions presented at 5 visual contrast levels (Figure 1B), ranging from 0 to 100%, with a static gray screen between trials. On half of the trials, we paired the visual stimulus with a 70 dB burst of white noise from a speaker positioned next to the screen (Figure 1C), affording 10 trials of each unique audiovisual stimulus condition (Figure 1C). Twelve recording sessions across six mice were spike sorted, and the responses of these sorted neurons were organized by trial type to compare across audiovisual stimulus conditions. We identified a total of 816 units across recordings, 161 (19.7%) of which were single units. Figure 1D-G demonstrates an example unit tuned for gratings aligned to the 30°-210° axis whose baseline and light-evoked firing rate are both increased by the sound.

**Figure 1.**
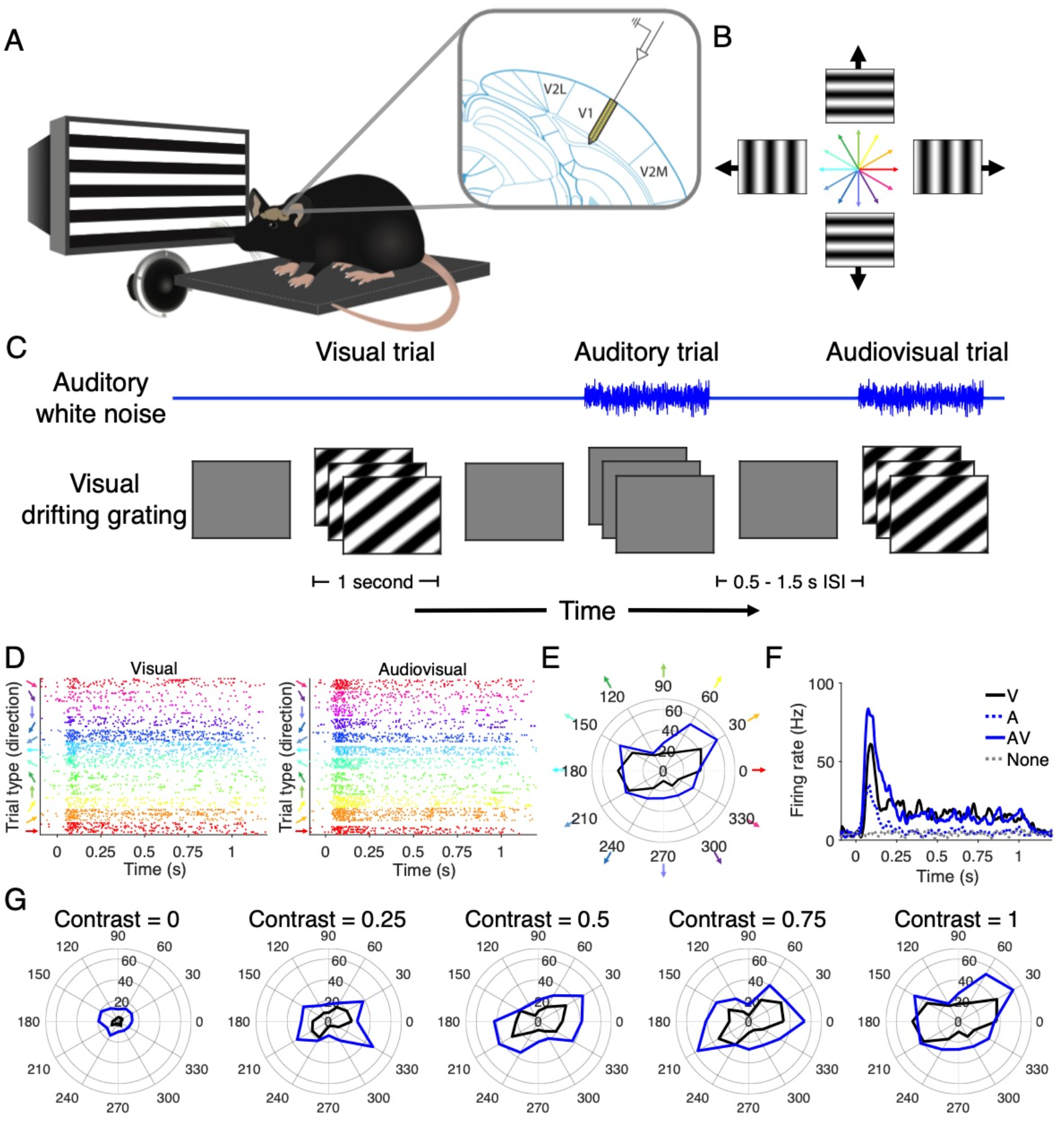
Audiovisual stimulus presentation. (A) Diagram (left) demonstrating that mice were head-fixed and presented with audiovisual stimuli from the right spatial field while electrophysiological recordings were performed in V1 (right). (B) Visual stimuli consisted of drifting gratings of 12 directions. (C) Auditory, visual, and audiovisual trials were randomly ordered and spaced with variable inter-stimulus intervals. (D) Raster plots of visual (left) and audiovisual (right) trials of an example neuron exhibiting visual orientation tuning. (E) Polar plot demonstrating the orientation tuning and magnitude of response (Hz) of the same example neuron in *E*. (F) PSTH of the same neuron in *E* demonstrating enhanced firing in response to audiovisual stimuli compared to unimodal stimuli. (G) Example neuron in *E* displays enhanced firing rate with sound across visual contrast levels.

Sound modulated the activity of the majority of V1 neurons. We used a generalized linear model (GLM) to classify neurons as light-responsive and/or sound-responsive based on their firing rate at the onset (0-300 ms) of each trial. We chose to classify neurons based on their onset response because the first 300 ms had the highest mutual information with both the visual contrast level as well as the drifting grating orientation (Figure 2A-C; Table 1). Using this classification method, we found that 86.2% (703/816) of units were responsive to increasing visual stimulus contrast levels, and of these visually responsive units, 80.1% (563/703 neurons, 12 recording sessions in 6 mice) were significantly modulated by the presence of sound (Figure 3A). Because the depth electrode penetrated all layers of V1, we were able to estimate the depth of each unit based on the amplitude of the spike waveform recorded by local electrodes. Surprisingly, we found that the majority of units across each depth were either sound-responsive or sound-modulated light-responsive (Figure 3F-H). We then constructed an average PSTH from the response profiles of sound-modulated light-responsive neurons, which revealed that the largest change in light-evoked firing rate occurred at the onset of the stimulus (Figure 3B). Averaged across neurons, we found a robust increase in the magnitude of the visually evoked response across visual contrast levels (Figure 3C; p(vis)=1.2e-100, p(aud)=1.6e-88, p(interact)=5.7e-4, paired 2-way ANOVA; p_c=0_=2.1e-51, p_c=0.25_=2.6e-62, p_c=0.5_=5.7e-75, p_c=0.75_=1.1e-81, p_c=1_=2.0e-81, post hoc Bonferroni-corrected paired t-test, Table 1). This difference was driven by the majority of neurons (95%) that increased their firing rate in the presence of sound. However, some neurons did exhibit lower light-evoked and sound-evoked firing rates relative to baseline.

**Figure 2.**
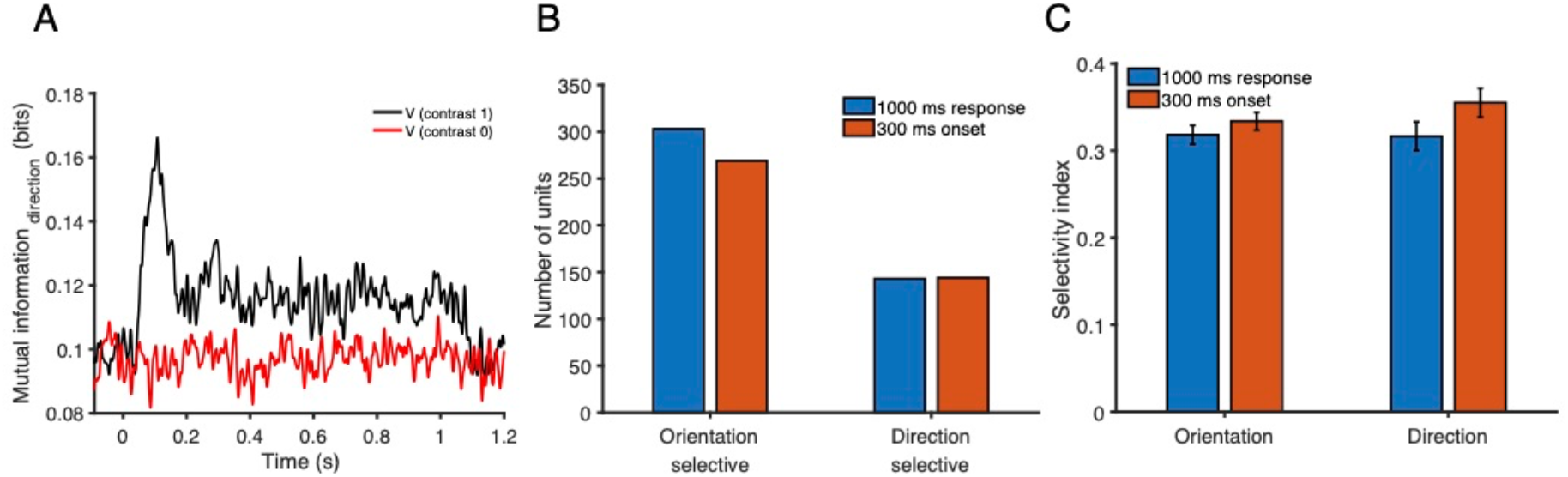
Classification based on sustained and onset responses. (A) Mutual information (MI) between neuronal responses and drifting grating direction, averaged across neurons. The black line is MI at full visual contrast, and the red line is MI at zero visual contrast. (B) We found a slight reduction in the number of neurons classified as orientation (269 vs 303 units) or direction selective (144 vs 143 units) when based on the initial 300 ms onset response compared to the entire 1000 ms response. (C) The OSI was similar (0.32 vs 0.33, n_1000_=303, _n=300_=269, p=0.30), as was the DSI (0.32 vs 0.36, n_1000_=143, _n=300_=144, p=0.10), of classified neurons when calculated using the initial 300 ms onset response compared the whole 1000 ms response.

**Figure 3.**
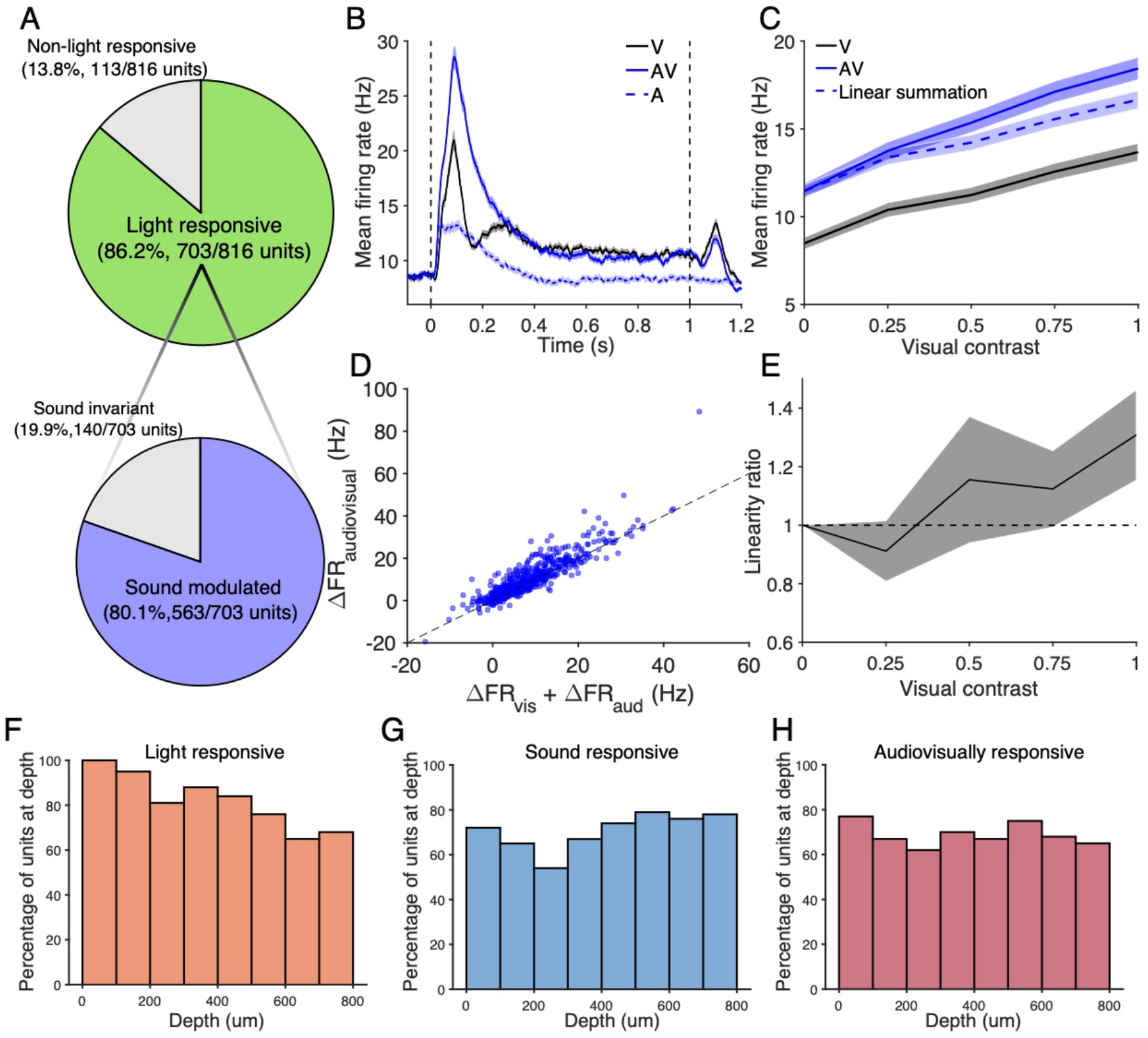
Sound enhances visual responses in a supra-linear manner. (A) Sound modulates visually evoked activity in 80.1% of light-responsive neurons in V1. (B) Comparison of visual, auditory, and audiovisual PSTHs averaged across all light-responsive sound-modulated neurons. Visual and audiovisual PSTHs correspond to the highest visual contrast level. (C) The magnitude of audiovisual onset responses (0-300ms) is greater than that of the visual response in light-responsive sound-modulated neurons (n=563, p(vis)=1.2e-100, p(aud)=1.6e-88, p(interact)=5.7e-4, 2-way repeated measures ANOVA; post hoc Bonferroni-corrected paired t-test). The expected linear sum of the unimodal auditory and visual responses is included. (D) At full visual contrast, the observed audiovisual response in the majority of neurons is greater than the linear sum of the unimodal auditory and visual responses. (E) A linearity ratio above 1 demonstrates audiovisual responses in V1 represent supra-linear integration of the unimodal signals (n=563, p=1.6e-12, Kruskal-Wallis test, post hoc Bonferroni-corrected Wilcoxon signed rank test). (F-H) Histograms demonstrating the percentage of neurons at each 100 um depth bin that were classified as light, sound, and audiovisually responsive, based on the recording electrode with the largest spike waveform amplitude.

This change in firing rate can be potentially supra-linear, linear or sub-linear based on whether the audiovisual response is, respectively, greater, equal or less than the sum of the unimodal light-evoked and sound-evoked firing rates. At medium to high visual contrast levels, integration of the audiovisual stimulus was predominantly supra-linear (Figure 3D-E; p=1.6e-12, Kruskal-Wallis test; p_c=0.25_=0.053, p_c=0.5_=0.004, p_c=0.75_=4.6e-8, p_c=1_=2.1e-5, post hoc Bonferroni-corrected Wilcoxon signed rank test, Table 1). In summary, these results show that sound supra-linearly increases the magnitude of the light-evoked response in the majority of V1 neurons.

### Sound reduces the orientation- and direction-selectivity of tuned neurons

Having observed sound-induced changes in the magnitude of the visual response, we next assessed how these changes in magnitude affected neuronal tuning profiles in the awake brain. Mouse V1 neurons typically have receptive fields tuned to a specific visual stimulus orientation and, to a lesser extent, stimulus direction (Métin et al, 1988; Rochefort et al., 2011; Fahey et al., 2019). To characterize these tuning profiles, we calculated orientation and direction-selective indices (OSI and DSI) in audiovisually responsive neurons. In addition to this magnitude-based metric, we also calculated pseudo-indices based on randomly shuffled permutations of neurons’ responses on each trial of orthogonal or opposite directions. Comparison of each neuron’s true OSI or DSI to its respective distribution of pseudo-indices allowed us to incorporate trial-wise variability into the selectivity criteria. We classified each neuron as orientation- or direction-selective whose true OSI or DSI, respectively, were greater than the 75^th^ percentile of its pseudo-indices. Figure 4A and 4C demonstrate the distribution of audiovisually responsive neurons’ selectivity indices, with additional shading indicating multi and single units. Figure 4B and 4D show example units along with the relationship between their true OSI or DSI and the distribution of pseudo-indices. Using this stringent selection criterion, we found that 47.8% (269/563) of neurons were orientation-selective, whereas 25.6% (144/563) were direction-selective. Surprisingly, we found a small reduction in the OSI from the visual to audiovisual conditions (Figure 4E; p=2.4e-6, paired Student’s t-test), which may reflect disproportionate changes in firing rate at the preferred versus orthogonal directions. We also found a slight reduction in DSI in the presence of sound (Figure 4F; p=6.4e-4, paired Student’s t-test). These sound-induced reductions in OSI and DSI were not as strong within single units (Figure 4E,F; p_OSI,single_=0.055, p_DSI,single_=0.033), and were relatively uniform across unit depth (Figure 4G). We also observed little shift in the preferred direction from the visual to audiovisual condition (Figure 4H), as calculated as half the complex phase of the response profile at full visual contrast (Niell and Stryker, 2008). In order to determine how sound affected the shape of the tuning profile, we aligned neurons’ tuning curves and normalized by each neuron’s response to the full contrast visual stimulus. Surprisingly, we found that sound enhanced responses across tuning bandwidth, an effect that was present across visual contrast levels (Figure 4I). Taken together, we observed that sound’s enhancing effect on visual response magnitude resulted in a mild reduction in tuning selectivity in orientation and direction-selective neurons.

**Figure 4.**
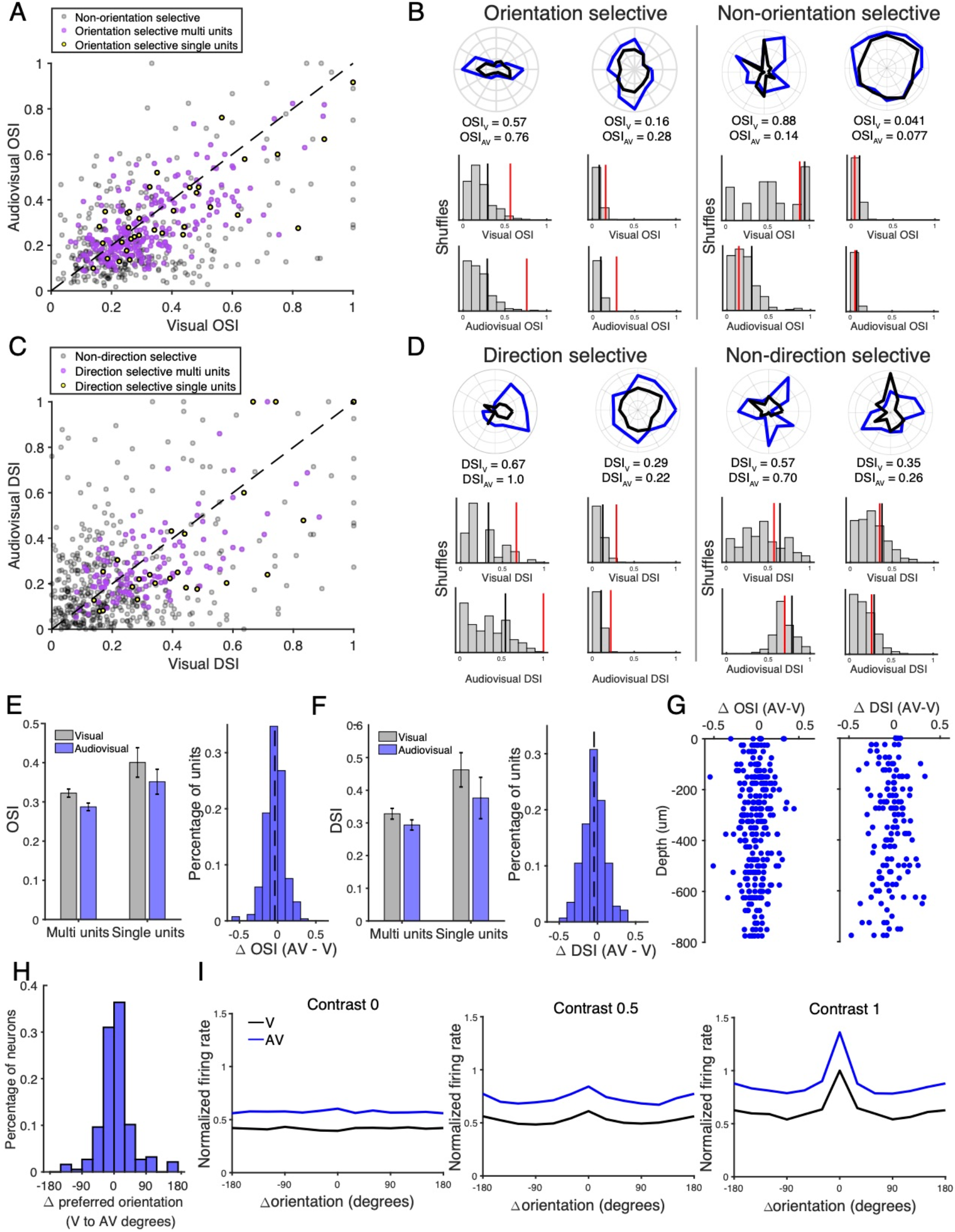
Sound reduces orientation- and direction-selectivity in tuned neurons. (A) Distribution of orientation selectivity index across audiovisually responsive units. Neurons with true OSI values above 75% of pseudo indices are shaded in purple (multi-units) or yellow (single units). (B) Example units from A, demonstrating each unit’s range of pseudo-indices (gray histogram), the true OSI value (red bar), and the 75% threshold (black bar). (C) Distribution of direction selectivity index across audiovisually responsive units. Again, red neurons have true DSI above 75% of pseudo indices. (D) Example units from C, again with DSI pseudo-index distributions (gray histogram), true DSI value (red bar), and 75% threshold (black bar). (E) A mild reduction in OSI was observed in multi-units (n=234, p=1.8e-5, paired t-test), with a trend in single units (n= 34, p=0.055, paired t-test). The distribution in ΔOSI shown on the right. (F) A mild reduction in DSI was observed in multi-units (n=120, p=7.4e-3, paired t-test) and in single units (n= 24, p=0.033, paired t-test). The distribution in ΔDSI shown on the right. (G) Change in OSI (left) and DSI (right) was relatively uniform across cortical depth. (H) Histogram depicting changes in preferred drifting grating directions, calculated using half of the complex phase, with sound in orientation-selective neuron. (I) Tuning curves under the visual and audiovisual conditions across visual contrast levels, averaged across neurons, show nonspecific sound-induced enhancement.

### Changes in neuronal response latency, onset duration, and variability in audiovisual compared to visual conditions

Behaviorally, certain cross-modal stimuli elicit shorter reaction times than their unimodal counterparts (Diederich and Colonius, 2004; Colonius and Diederich, 2017; Meijer et al., 2018). Therefore, we hypothesized that sound reduces the latency of the light-evoked response at a neuronal level as well. For each neuron, we calculated the response latency as the first time bin after stimulus onset at which the firing rate exceeded 1 standard deviation above baseline (Figure 5A), and found that sound reduced the response latency across contrast levels (Figure 5B; p(vis)=6.9e-4, p(aud)=6.8e-15, p(interact)=0.045, paired 2-way ANOVA; p_c=0.25_=2.3e-4, p_c=0.5_=7.1e-12, p_c=0.75_=4.6e-5, p_c=1_=9.9e-4, post hoc Bonferroni-corrected paired t-test, Table 1). We additionally calculated the slope of the onset response of light-responsive sound-modulated neurons, measured from trial onset until the time at which each neuron achieved its peak firing rate (Figure 5C). We found that sound increased the slope of the onset response (Figure 5D; p(vis)=3.5e-121, p(aud)=2.7e-15, p(interact)=0.038, paired 2-way ANOVA; p_c=0.25_=1.4e-4, p_c=0.5_=8.9e-13, p_c=0.75_=3.6e-12, p_c=1_=5.5e-8, post hoc Bonferroni-corrected paired t-test, Table 1), both indicating that the response latency was reduced in the audiovisual condition compared to the visual condition. Additionally, the duration of the light-evoked response, defined as the full width at half maximum of the peak onset firing rate, increased in the presence of sound (Figure 5E,F; p(vis)=1.3e-10, p(aud)=8.7e-98, p(interact)=0.23, paired 2-way ANOVA). Both of these timing effects were preserved across contrast levels. Therefore, the latency and onset duration of neuronal audiovisual responses of V1 neurons is enhanced compared to visual responses.

**Figure 5.**
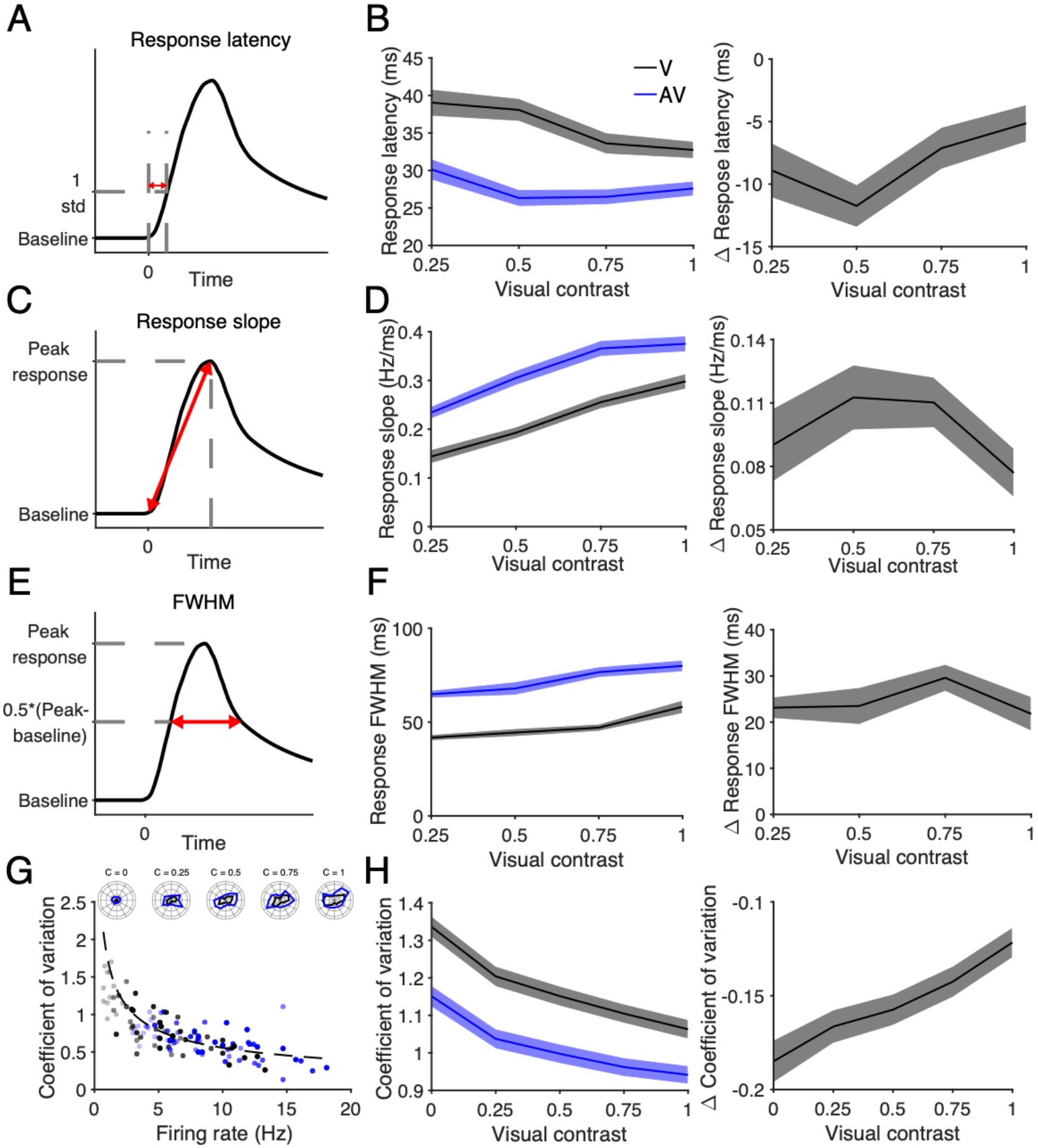
Changes in neuronal response latency, onset duration, and variability in audiovisual compared to visual conditions. (A) Diagram of the calculation of response latency, the first time bin in which the FR exceeds 1 std above baseline. (B) Audiovisual response latency is less than that of the visual response (left: absolute, right: difference; p(vis)=6.9e-4, p(aud)=6.8e-15, p(interact)=0.045, paired 2-way ANOVA, post hoc Bonferroni-corrected paired t-test, Table 1). (C) Diagram of the calculation of response onset slope, the peak change in FR over the latency to peak response. (D) The slope of the audiovisual response is greater than that of the visual response (left: absolute, right: difference; n=563, p(vis)=3.5e-121, p(aud)=2.7e-15, p(interact)=0.038, paired 2-way ANOVA, post hoc Bonferroni-corrected paired t-test). (E) Diagram of the calculation of FWHM, the width of the onset response at half maximum FR. (F) The FWHM of the audiovisual response is greater than that of the visual response (left: absolute, right: difference; n=367, p(vis)=1.3e-10, p(aud)=8.7e-98, p(interact)=0.23 paired 2-way ANOVA). (G) An example neuron demonstrating that increased response magnitude corresponds to lower CV according to an inverse square root relationship. The black and blue dots represent visual and audiovisual responses, respectively, and the dot transparency corresponds to visual contrast level. The dotted lines are fitted y=c/sqrt(x) curves, where c is a constant. The above inset is the polar plots corresponding to the example neuron. (H) Lower coefficient of variation indicates reduced response variability in audiovisual compared to visual responses (left: absolute, right: difference; n=563, p(vis)=0.28, p(aud)=4.2e-103, p(interact)=0.38, paired 2-way ANOVA).

Having observed changes in response magnitude and timing, we next investigated the effect of sound on the variability of light-evoked responses. If individual neurons encode the visual stimulus using changes in their firing rate, a more consistent response would entail less spread in the response magnitude relative to the mean response across trials of a single stimulus type. We quantified this relationship using the coefficient of variation (CV) defined as the ratio of the standard deviation to the response mean (Gur et al., 1997). We hypothesized that sound reduces the CV of light-evoked responses, corresponding to reduced response variability and higher signal-to-noise ratio. Figure 5G depicts the relationship between response magnitude and CV in an example sound-modulated light-responsive neuron, demonstrating that increased response magnitude correlates with reduced CV. Consistent with sound increasing the visual response magnitude in the majority of sound-modulated light-responsive neurons (Figure 3), we observed a reduction of CV in the audiovisual condition relative to the visual condition when averaged across these neurons (Figure 5H; p(vis)=0.28, p(aud)=4.2e-103, p(interact)=0.38, paired 2-way ANOVA). Taken together, these results indicate that sound not only modulates the magnitude of the visual response (Figure 3), but also improves the timing and consistency of individual neurons’ responses (Figure 5).

### Sound-induced movement does not account for sound’s effect on visual responses

It is known that whisking and locomotive behaviors modulate neuronal activity in mouse visual cortex (Niell and Stryker, 2010; Mesik et al., 2019) and auditory cortex (Nelson et al., 2013; Schneider and Mooney, 2018; Bigelow et al., 2019). Therefore, having established that sound robustly modulates visual responses (Figure 3), we tested whether and to what extent these observed changes were more accurately attributable to sound-associated uninstructed movement in the mouse subjects. In an additional cohort of mice, we performed V1 extracellular recordings with the same audiovisual stimuli described above while recording movement activity of the mice throughout stimulus presentation (Figure 6A). Despite being head-fixed to afford stable electrophysiological recordings, the mice were positioned on a smooth stage that freely allowed volitional movements. We used the publicly available Facemap software to process the video data (Stringer et al., 2019). Using its pre-programmed GUI, regions of interest (ROI) were placed around the whiskers and face of mouse subjects in order to identify and quantify whisking and facial behavior (Figure 6B, bottom). ROIs were also placed on the limbs to identify locomotion (Figure 6B, middle), and additional ROIs were distributed across the entire mouse subject, including the face and limbs, in order to capture general nonspecific movements (Figure 6B, top).

**Figure 6.**
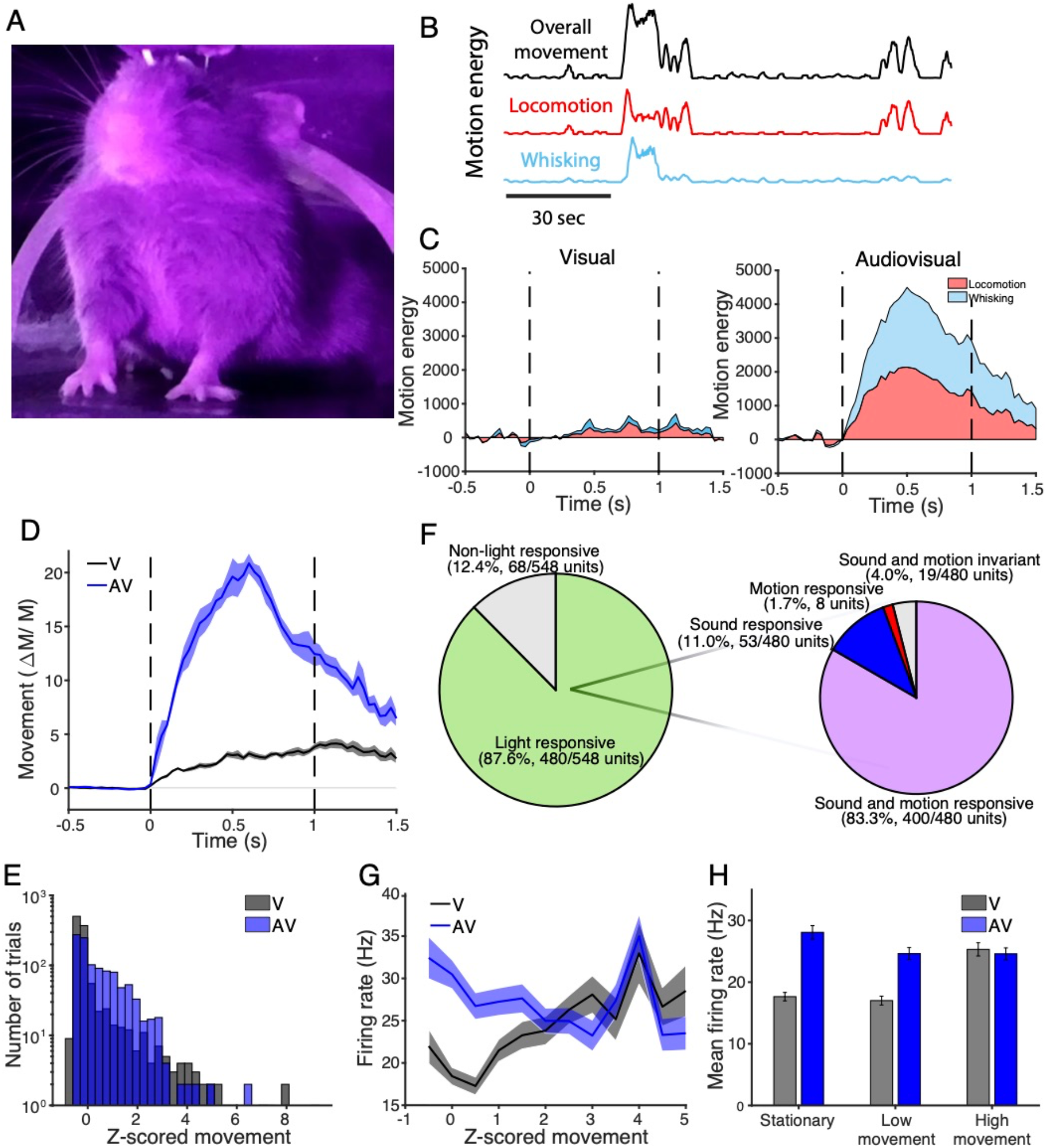
Sound modulates visual activity when controlling for stimulus-induced movement. (A) A still image demonstrating the video capture of mouse subjects during recording sessions. (B) Sample whisking, locomotion, and overall movement trace outputs using Facemap video analysis. (C) Average stimulus-aligned locomotion and whisking behavior on visual (left) and audiovisual (right) trials, relative to baseline 100ms prior to stimulus onset. (D) Mice displayed more general movement, including both locomotion and whisking, response to audiovisual trials than in visual trials (n=9 recording sessions; p=4.0e-7, paired t-test). (E) Histogram of trials’ z-scored movements show a range of levels of movement during both visual and audiovisual trials. (F) Venn diagram demonstrating that 96% of light-responsive neurons exhibited some combination of sound- and movement-responsiveness. (G) Comparison of firing rate of sound- and motion-modulated light-responsive neurons across trials with a range of z-scored movement. (H) Responses to audiovisual stimuli evoke larger magnitude responses than visual stimuli when mice were stationary (z-score<-0.5) or displayed low to moderate movement (−0.5<z-score<1.5), but responses were not significantly different when mice displayed the highest amount of movement (z-score>1.5; p(motion)=1.6e-8, p(aud)=1.6e-8, p(interact)=3.1e-8, 2-way ANOVA, post hoc Bonferroni-corrected two-sample t-test).

The energy output of each of these ROI regions throughout the video recording was then aligned with the audiovisual stimulus in order to process, identify, and quantify stimulus-correlated movements. For each recording session, we calculated averages for each trial type in order to compare visual and audiovisual movement responses for each mouse subject. We found that both visual and auditory stimuli did evoke whisking and locomotive behavior in mice, with combined audiovisual stimuli evoking a larger degree of both behaviors than isolated visual stimuli (Figure 6C). Using the general movement trace, which included both locomotive and whisking behavior, we subtracted the 100ms baseline prior to trial onset from the movement trace throughout the trial, and then divided by that baseline value in order to calculate fold increase over baseline. Using this method, we found that movement was higher during audiovisual trials compared to visual trials (Figure 6D; p=4.0e-7, paired t-test). However, there were many visual trials in which substantial movement occurred, as well as audiovisual trials in which little movement was detected (Figure 6E). Because of this variability in sound-induced movement, we were able to control for movement when comparing visual and audiovisual activity in the recorded neurons.

We used a GLM to classify each neuron as light-, sound-, and/or motion-responsive based on the neuron’s firing rate and mouse’s general movement activity during the onset (0-300ms) of the trial. The vast majority of light-responsive neurons, 83.3% (400/480), displayed both sound- and motion-modulated visual responses (Figure 6F). 11.0% (53/480) and 1.7% (8/480) of light-responsive neurons were purely sound- or motion-modulated, respectively. An additional 4.0% (19/480) were invariant to sound or motion. We then compared the visually and audiovisually evoked firing rates of neurons when accounting for movement. Among sound- and motion-modulated light-responsive neurons, the firing rate was higher on audiovisual trials than visual trials when movement was held constant (Figure 6G), especially when mice showed limited movement. On trials in which the mice were largely stationary (z-score<-0.5, 49% of visual trials, 33% of audiovisual trials) or displayed moderate levels of movement (−0.5<z-score<1.5, 45% of visual trials, 55% of audiovisual trials), the mean firing rate of neurons was 54-62% higher when sound was presented than when sound was absent. The firing rates under the two stimulus conditions converged on trials in which the mice displayed high movement activity (z-score>1.5, 4.9% of visual trials, 12% of audiovisual trials; Figure 6G,H; p(move)=1.6e-8, p(aud)=1.6e-8, p(interact)=3.1e-8, unbalanced 2-way ANOVA; p_stationary_=1.0e-15, p_low motion_=3.1e-10, p_high motion_=0.59, post hoc Bonferroni-corrected two-sample t-test, Table 1). Notably, increasing movement activity was correlated with increased firing rates on visual trials, but was correlated with decreasing firing rates among audiovisual trials (Figure 6H). These results indicate that sound modulated visually evoked neuronal activity even when accounting for sound-induced movement in awake mice, with the exception of when mice display high amount of movement, during which there was little effect of sound on firing rates.

### Sound and movement have distinct and complementary effects on visual responses

To further parse out the role of sound and movement on audiovisual responses, we used a separate GLM to capture the time course of these parameters’ effects on visually evoked activity. For each neuron, we used a GLM with a sliding 10ms window to reconstruct the PSTH based on the visual contrast level, sound presence, and general movement, which included both locomotion and whisking behavior, during that time window (Figure 7A). Figure 7B shows two example neurons in which the GLM estimated the light-evoked, sound-evoked, and audiovisually evoked PSTHs using the average movement for each trial type. Across neurons, the GLM-estimated PSTHs accurately reconstructed the observed PSTHs (Figure 7C-E). We leveraged the coefficients fit to each neuron (Figure 7A) to estimate the unique contribution of each predictor to the firing rates as a function of time (see Materials and Methods). When the movement parameter was minimized, sound predominantly enhanced neuronal activity at the onset of the visual response and suppressed activity during the response’s sustained period (Figure 7F; n=295 fitted neurons, paired t-test at each time window [1391], α=3.6e-5). Conversely, movement had limited effect on the onset activity in the absence of sound, but rather primarily enhanced firing rates during the response’s sustained period (Figure 7G, red trace; n=295 fitted neurons, paired t-test at each time window [1391], α=3.6e-5). In a separate analysis, the nonspecific movement variable was replaced by two independent variables representing locomotion and whisking, and a similar GLM including coefficients for each movement subtype was fit to each neuron’s PSTH. The estimated visual response PSTH with average locomotion and minimal whisking, as well as with average whisking and minimal locomotion, are also on display in Figure 7G (teal and green traces, respectively). Together, sound and movement had complementary effects in which both the onset and sustained portions of the visual response were enhanced compared to the isolated visual response (Figure 7H, pink trace; n=295 fitted neurons, paired t-test at each time window [1391], α=3.6e-5). Again notably, the peak onset response under the audiovisual condition was lower when movement was included in the estimate (Figure 7H, pink versus blue traces). We additionally grouped neurons by their cortical depth, and found that the distinct effects that sound and movement had on visual responses were largely preserved across layers (Figure 7I,J), although were slightly larger in magnitude at superficial depths. These findings indicate not only that movement is unable to account for the changes in onset response reported above, but also that sound and motion have distinct and complementary effects on the time course of visually evoked activity in V1.

**Figure 7.**
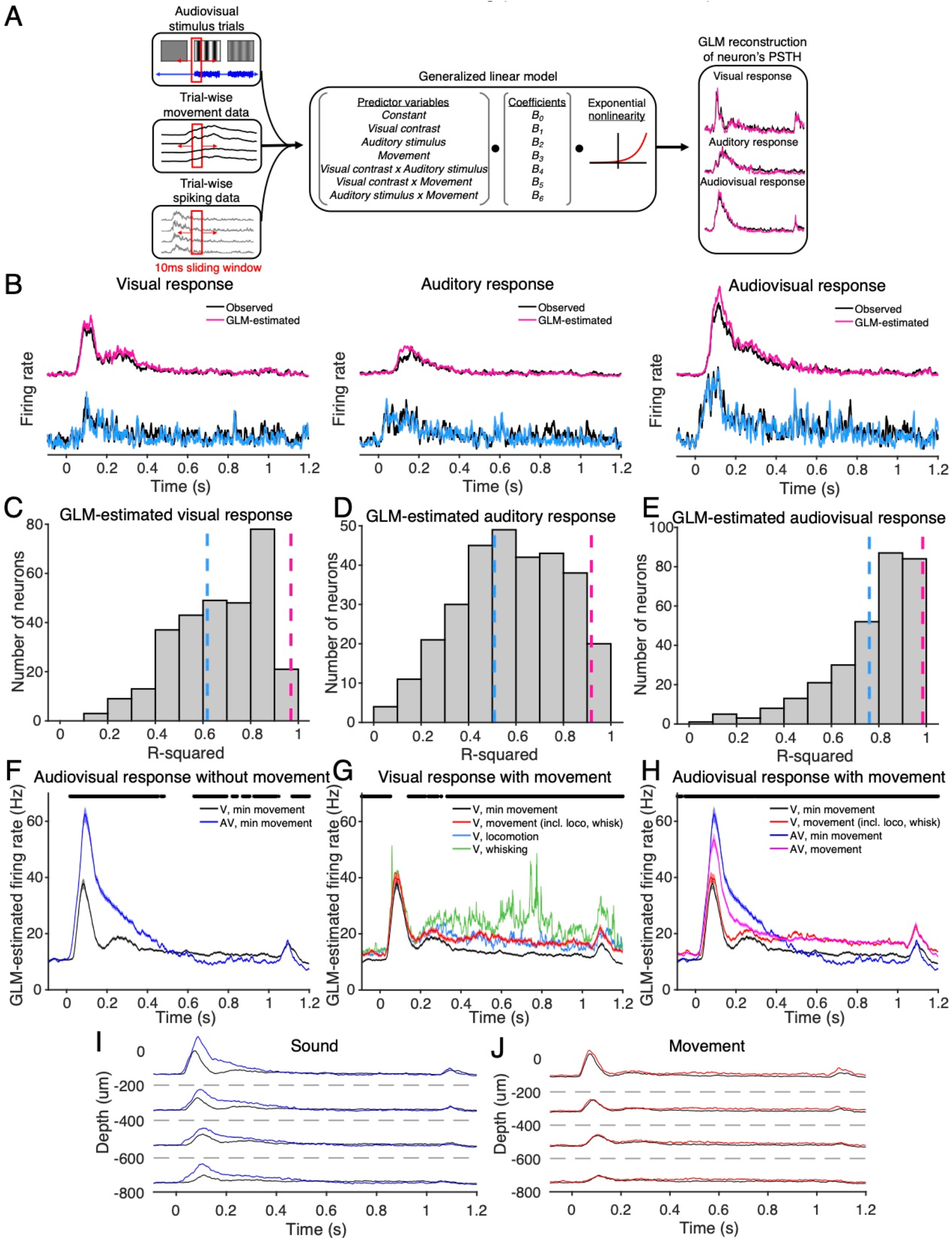
Sound and movement modulate visual responses in distinct but complementary ways. (A) Diagram illustrating the use of a GLM to reconstruct individual neurons’ PSTHs based on neuronal responses and mouse movement during stimulus presentation. The GLM was then used to predict the time course of neuronal responses audiovisual stimuli with and without movement. (B) Observed trial-averaged PSTHs for visual-only (left), auditory-only (middle), and audiovisual (right) trials overlaid with GLM estimates based on the selected stimulus features for two example units (blue and pink). (C-E) Histograms demonstrating R^2^ values of the GLM-estimated PSTHs, averaged across sound- and motion-modulated light-responsive neurons. Moderate to high R^2^ values across the population indicate a good ability for the GLM to estimate neuronal firing rates. The dashed pink and blue line shows the R^2^ value associated with the two example units in B. (F-H) GLM-predicted visually evoked PSTHs with and without sound and motion. Asterisks indicate time windows in which there was a significant difference between the *light* prediction and the *light+sound, light+motion*, and *light+sound+motion* predictions, respectively. (F) Excluding motion highlights that sound primarily enhances the onset response. Asterisks indicate time windows in which there was a significant difference (n=343 fitted neurons; paired t-test, α=3.6e-5). (G) Excluding sound highlights that nonspecific motion (red), as well as locomotion (blue) and whisking (green), primarily enhance the sustained portion of the response. Asterisks indicate time windows in which there was a significant difference (n=343 fitted neurons; paired t-test, α=3.6e-5). (H) Sound and motion together enhance both the onset and sustained periods of the visually evoked response compared to the isolated visual response. (n=343 fitted neurons; paired t-test, α=3.6e-5). (I) Sound’s enhancing effect on the visual response onset was slightly stronger at superficial cortical depths compared to deeper layers, when averaged across neurons in 200um depth bins. (J) Similarly, movement’s enhancing effect on the visual response sustained portion was slightly stronger at superficial cortical depths, when averaged across neurons in 200um depth bins.

### Decoding of the visual stimulus from individual neurons is improved with sound

Behaviorally, sound can improve the detection and discriminability of visual responses, however whether that improved visual acuity is reflected in V1 audiovisual responses is unknown. Many studies have reported how sound affects visual responses in V1, but whether these changes improve neuronal encoding of the visual stimulus in the awake brain has not been robustly demonstrated. The increase in response magnitude and decrease in CV suggest that sound may improve visual stimulus discriminability in individual V1 neurons. Consistent with these changes in response magnitude and variability, we observed sound-induced improvements in the *d’* sensitivity index between responses to low contrast drifting grating directions among orientation- and direction-selective neurons (Figure 8A,B), further indicating improved orientation and directional discriminability in individual neurons. To directly test this hypothesis, we used the neuronal responses of individual neurons to estimate the visual stimulus drifting grating orientation and direction. We trained a maximum likelihood estimate (MLE)-based decoder (Montijn et al., 2014; Meijer et al., 2017) on trials from the preferred and orthogonal orientations in orientation-selective neurons and on trials from the preferred and anti-preferred directions in direction-selective neurons. We used leave-one-out cross-validation and cycled the probe trial through the repeated trials of the stimulus condition in order to calculate the mean decoding performance. The MLE decoder’s output was the orientation or direction with the maximum posterior likelihood based on the training data and probe trial (Figure 8C). This decoding technique achieves high decoding accuracy (Figure 8D). When averaged across sound-modulated orientation-selective neurons, decoding performance was improved on audiovisual trials compared to visual trials (Figure 8E; p(vis)=4.8e-112, p(aud)=1.7e-11, p(interact)=1.0e-5, paired 2-way ANOVA; p_c=0_=0.78, p_c=0.25_=1.5e-4, p_c=0.5_ =2.2e-11, p_c=0.75_ =0.21, p_c=1_=1.4e-6), with the greatest improvements at low to intermediate contrast levels. We applied this approach to sound-modulated direction-selective units and found similar sound-induced improvements in decoding accuracy (Figure 8G; p(vis)=1.2e-15, p(aud)=6.9e-4, p(interact)=0.82, paired 2-way ANOVA). Furthermore, similar effects were observed in both single and multi-units (Figure 8F,H). These results demonstrate that sound-induced changes in response magnitude and consistency interact in order to improve neuronal representation of the visual stimulus in individual neurons.

**Figure 8.**
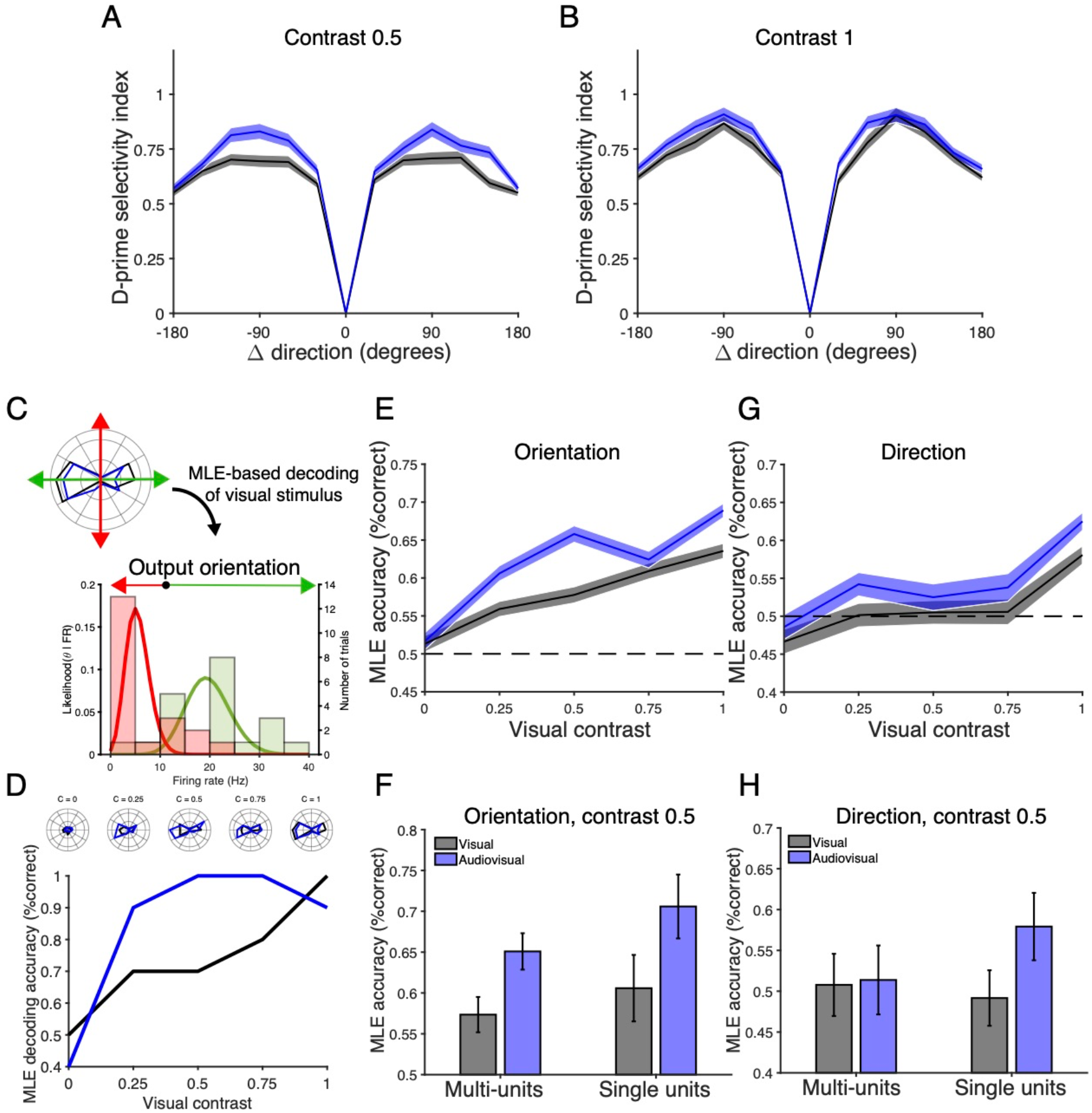
Sound improves decoding of drifting grating direction and orientation in individual neurons. (A-B) The *d’* sensitivity index between neuronal responses to drifting grating directions, averaged across orientation- and direction-selective neurons. Enhancements are observed at low visual contrast (A), whereas minimal changes are present at full contrast (B). (C) Diagram illustrating MLE-based decoding of an individual neuron’s preferred versus orthogonal orientations. (D) Performance of the MLE decoder, trained on an example orientation-selective neuron, in decoding the neuron’s preferred versus orthogonal orientations. The neuron’s polar plots are shows in the above inset. (E) Absolute difference in decoding accuracy of preferred versus orthogonal orientations, averaged across sound-modulated orientation-selective neurons, demonstrating higher performance in the audiovisual condition (n=269, p(vis)=4.8e-112, p(aud)=1.7e-11, p(interact)=1.0e-5, paired 2-way ANOVA). (F) Similar improvements in decoding were observed in both multi- and single units. (G) Absolute difference in decoding accuracy of preferred versus anti-preferred directions, averaged across sound-modulated direction-selective neurons, again with improved performance on audiovisual trials compared to visual trials (n=144, p(vis)=1.2e-15, p(aud)=6.9e-4, p(interact)=0.82, paired 2-way ANOVA).(H) The improvement in decoding probe trial direction was principally driven by single units, with limited effects observed in multi-units.

### Population-based decoding of the visual stimulus improves with sound

V1 uses population coding to relay information about the various stimulus dimensions to downstream visual areas (Montijn et al., 2014, Berens et al., 2012), so we next tested whether these improvements in visual stimulus encoding in individual neurons extended to the population level. We again used a leave-one-out cross-validation approach when training and testing the decoder (Figure 9A). Unsurprisingly, decoding accuracy improved as more neurons were included in the population (Figure 9B). We began by using the MLE-based decoder to perform pairwise classification of visual drifting grating directions based on neuronal population activity. At full visual contrast, there was little difference between the performance on visual and audiovisual trials. However, at low to intermediate visual contrast levels, classification performance increased on audiovisual trials as compared to visual trials (Figure 9C). This improvement in performance was greatest when comparing orthogonal drifting grating orientations (Figure 9D; p(vis)=2.6e-98, p(aud)=0.098, p(interact) = 1.7e-82, 2-way ANOVA; p_c=0_=2.1e-7, p_c=0.25_=1.5e-12, p_c=0.5_,=4.1e-9, p_c=0.75_=1.4e-4; p_c=1_=9.4e-4, post hoc Bonferroni-corrected paired t-test, Table 1). However, there was limited sound-induced improvement in decoding opposite drifting grating directions (Figure 9E, p(vis)=4.2e-90, p(aud)=0.87, p(interact)=8.1e-6, 2-way ANOVA; p_c=0_=0.21, p_c=0.25_=5.9e-4, p_c=0.5_=2.5e-4, p_c=0.75_=0.48, p_c=1_=0.97, post hoc Bonferroni-corrected paired t-test, Table 1).

**Figure 9.**
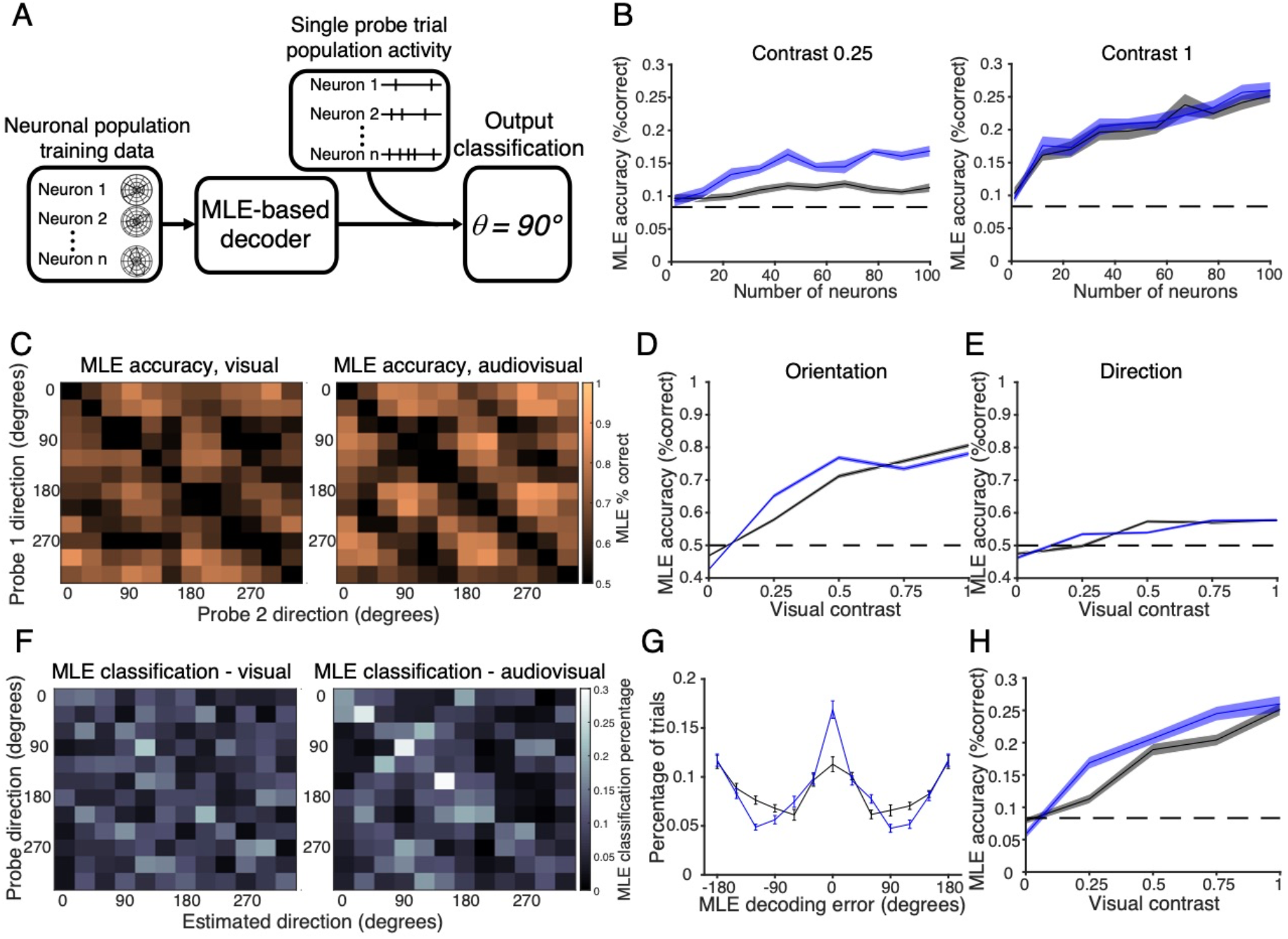
Sound improves accuracy of population-based visual stimulus decoding. (A) Schematic illustrating the decoding of the drifting grating direction using an MLE decoder trained on neuronal population activity. (B) Accuracy of MLE decoding 1-of-12 drifting grating options improved as the neuronal population size included in the decoder increases. Visual contrast 0.25 is on the left, and full visual contrast is on the right. (C) Accuracy of MLE pairwise classification of drifting gratings on visual (left) and audiovisual (right) trials, contrast 0.25. (D) MLE decoding accuracy when classifying orthogonal drifting grating orientations improved with sound (n=50 randomizations, p(vis)=2.6e-98, p(aud)=0.098, p(interact)=1.7e-82, 2-way ANOVA, post hoc Bonferroni-corrected paired t-test). (E) MLE decoding accuracy when classifying opposite drifting grating directions, demonstrating limited effect of sound on performance (n=50 randomizations, p(vis)=4.2e-90, p(aud)=0.87, p(interact)=8.1e-6, 2-way ANOVA, post hoc Bonferroni-corrected paired t-test). (F) Heat map of actual vs MLE-output directions under visual (left) and audiovisual (right) trials, contrast 0.25. MLE decoder could choose between all 12 drifting grating directions. (G) MLE decoder classification error, comparing estimated direction to actual direction. (H) Overall decoding accuracy of MLE decoder when choosing between all 12 drifting grating directions improved with sound (n=10 randomizations, p(vis)=8.7e-55, p(aud)=4.2e-4, p(interact)=3.3e-4, 2-way ANOVA, post hoc Bonferroni-corrected paired t-test).

Expanding on the pairwise discriminability approach, the MLE-based decoder allowed us to also perform classification of 1 out of all 12 drifting grating directions. When trained and tested in this fashion, MLE decoding performance again improved at low to intermediate contrast levels on audiovisual trials (Figure 9F-H), before reaching asymptotic performance at full visual contrast (Figure 9H; p(vis)=8.7e-55, p(aud)=4.2e-4, p(interact)=3.3e-4, 2-way ANOVA; p_c=0_=0.011, p_c=0.25_=2.9e-4, p_c=0.5_=0.090, p_c=0.75_=0.0054, p_c=1_=0.57, post hoc Bonferroni-corrected paired t-test, Table 1). Similar results were found when organizing the neurons by recording session instead of pooling all neurons together (data not shown). Taken together, these results indicate that sound improves neuronal encoding of the visual stimulus both in individual neurons and at a population level, especially at intermediate visual contrast levels.

### Sound improves stimulus decoding when controlling for sound-induced movements

It is known that sensorimotor inputs shape V1 visual responses (Niell and Stryker, 2010; Mesik et al., 2019), and locomotion improves visual processing in V1 (Dadarlat and Stryker, 2017). So we next tested whether the observed sound-induced improvement in visual stimulus representation (Figure 8, 9) was attributable to sound’s effect on visual responses or indirectly via sound-induced movement. As we previously observed, sound and movement enhanced the onset and sustained portion of the visual response, respectively (Figure 7). We therefore hypothesized that the improvement on MLE decoding performance, based on the visual response onset, would be present even when accounting for sound-induced uninstructed movements. We tested this hypothesis by expanding on the GLM-based classification of neurons described in Figure 7. Using the same GLM generated for each neuron, we independently modified either the sound or movement variables and their associated pairwise predictors to their lowest values, and then used the GLM coefficients and the exponential nonlinearity to estimate each neuron’s audiovisual response magnitude (Figure 10A, Materials and Methods). We then input these estimated trial-wise neuronal responses into the same MLE-based decoder described above (Figure 8,9). Using this approach, we found that in individual orientation-selective neurons, controlling for the effect of motion on audiovisual trials had little effect on the improvement in decoding accuracy on audiovisual trials (Figure 10B-C; p(vis)=6.2e-81, p(aud)=5.3e-3, p(interact)=0.15, paired 2-way ANOVA, Table 1). However, instead, regressing out sound from the audiovisual responses resulted in decoding accuracy that more closely resembled that of visual trials (Figure 10B-C; p(vis)=7.7e-72, p(aud) = 0.23, p(interact)=0.11, paired 2-way ANOVA, Table 1). These results in individual neurons suggest that sound and not movement primarily drives the improvement in decoding accuracy on audiovisual trials. We found similar results when implementing this approach in the MLE-based population decoder. We again found that that decoding performance on audiovisual trials when regressing out motion was still significantly improved compared to that on visual trials (Figure 10D-E; p(vis)=2.5e-46, p(aud)=2.1e-13, p(interact)=9.3e-5, 2-way ANOVA; p_c=0_=0.94, p_c=0.25_=1.0e-4, p_c=0.5_=0.010, p_c=0.75_=0.0023, p_c=1_=0.021, Bonferroni-corrected paired t-test). Furthermore, alternatively regression out sound from audiovisual trials resulted in population decoding performance similar to that on visual trials (Figure 10D-E; p(vis)=3.3e-43, p(aud)=0.88, p(interact)=2.4e-6, 2-way ANOVA; p_c=0_=0.87, p_c=0.25_=0.039, p_c=0.5_=0.19, p_c=0.75_=0.080, p_c=1_=0.0025, Bonferroni-corrected paired t-test). These results demonstrate that sound improves visual stimulus decoding on audiovisual trials at both a single neuron and population level. Moreover, this enhancement persists when controlling for sound-induced motion.

**Figure 10.**
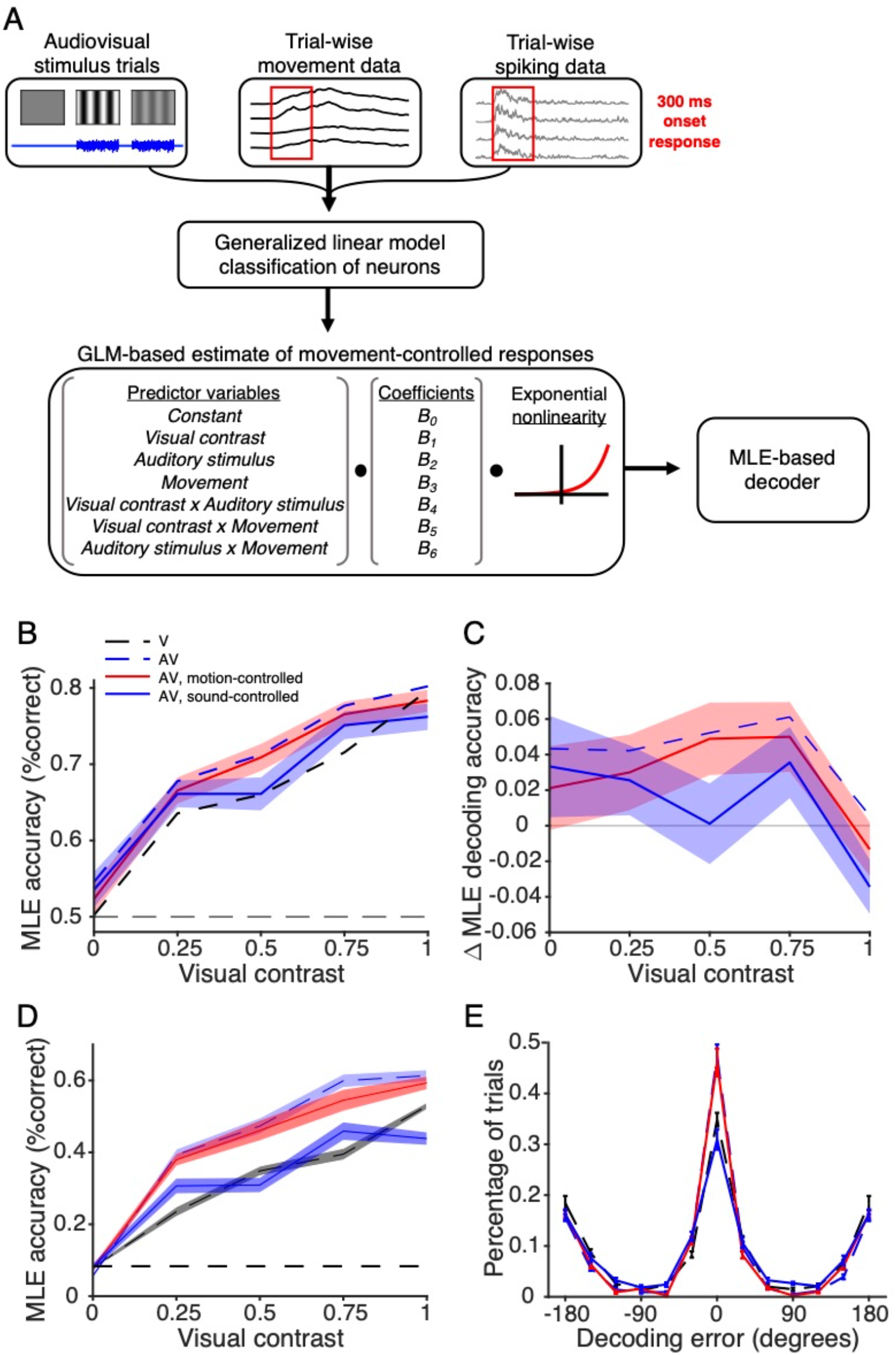
Sound improved decoding performance when controlling for motion. (A) Diagram illustrating the use of a GLM to calculate each predictor variable’s coefficient. These are then used when varying the predictor variables to estimate trial-wise neuronal responses, which are then input into the MLE-based decoder. (B) Absolute accuracy of decoding orientation among orientation-selective, sound/motion-modulated light-responsive neurons, comparing visual responses (black, dotted) to audiovisual responses (blue, dotted), audiovisual responses when regressing out motion (red, solid) and audiovisual responses when regressing out sound (blue, solid). (C) Relative decoding accuracy compared to decoding on visual trials. Regressing out motion still preserved improved performance compared to visual trials (n=90 neurons, p(vis)=6.2e-81, p(aud)=5.3e-3, p(interact)=0.15, paired 2-way ANOVA), whereas regressing out sound resulted in comparable performance to visual trials (n=90 neurons, p(vis)=7.7e-72, p(aud)=0.23, p(interact)=0.11, paired 2-way ANOVA). (D) Population decoding accuracy of population-based decoder on audiovisual trials (blue, dotted) is preserved even when controlling for motion (red, solid; n=10 randomizations, p(vis)=2.5e-46, p(aud)=2.1e-13, p(interact)=9.3e-5, 2-way ANOVA; Bonferroni-corrected paired t-test), whereas controlling for sound (blue, solid) resembles decoding performance on visual trials (black dotted; n=10 randomizations, p(vis)=3.3e-43, p(aud)=0.88, p(interact)=2.4e-6, 2-way ANOVA; Bonferroni-corrected paired t-test). (E) MLE decoder classification percentage, comparing estimated direction to actual direction, contrast 0.5. Little difference is observed between audiovisual trials and audiovisual trials when controlling for motion, whereas both are more accurate than visual trials.

## Discussion

Audiovisual integration is an essential aspect of sensory processing (Stein et al., 2020). In humans, audiovisual integration is used in everyday behaviors such as speech perception and object recognition (Fujisaki et al., 2014). The goal of the present study was to test whether sound drives improvement in encoding and decoding of visual stimuli in awake subjects, and to test the hypothesis that sound improves neuronal encoding of visual stimuli in V1 independent of sound-induced movement. We performed extracellular recordings in V1 while presenting combinations of visual drifting gratings and auditory white noise and recording movement of awake mice. The drifting gratings were presented at a range of visual contrast levels to determine the threshold levels at which sound is most effective. As in previous studies, we found neurons in V1 whose spontaneous and visually evoked firing rates are modulated by sound (Figure 3). Notably, the effects we observed were stronger and more skewed towards response-enhancing than in previous studies (80.3% of neurons were modulated by sound, with ∼95% exhibiting sound-induced increases in firing rate). When accounting for movement in awake animal subjects, we found that the neurons’ audiovisual responses represented a mixed effect of both sound- and movement-sensitivity (Figure 6), an effect in which sound primarily enhances the onset response whereas movement complementarily enhances the sustained response (Figure 7). We also found that the sound-induced changes in response magnitude and consistency combined to improve the discriminability of drifting grating orientation and direction in individual neurons (Figure 8) and at a population level (Figure 9). The improvements in neuronal encoding were most pronounced at low to intermediate visual contrast levels, a finding consistent with the current understanding that audiovisual integration is most beneficial for behavioral performance under ambiguous unisensory conditions (Gleiss and Kayser, 2012; Meijer et al., 2018; Stein et al., 2020), as found in human psychophysics (Lipper et al., 2007; Chen et al., 2011). Importantly, the improvement in neuronal encoding was based on firing at the onset of the visual response, indicating that the auditory signal itself is responsible for improvements in visual encoding and not attributable to uninstructed movements. This was directly demonstrated by the persistence of sound-induced improvements in stimulus decoding, even when controlling for the effect of motion (Figure 10).

### Auditory and locomotive inputs distinctly shape visual responses

We find that sound and movement have distinct and complementary effects on visual responses. Previous work found that locomotion modulates neuronal responses in the visual cortex in the presence of both sounds and visual stimuli but did not find an audio-specific interaction of locomotion’s effect (McClure and Polack, 2019). Our results have revealed this component possibly because of the dynamics evoked by our sound stimulus, a white burst at a moderate sound pressure level which elicited a partial locomotive response. In our analysis, stimulus decoding relied largely on neuronal responses during the stimulus onset period. Therefore, despite robust differences in movement during visual and audiovisual trials, motion, which affected neuronal responses over slower time scales, only partially contributed to these changes in decoding (Figure 10). Our focus on the onset response was based on our initial finding that mutual information between the neuronal responses and visual stimuli was highest during this onset period, a finding supported by previous studies (Figure 2; Dadarlat and Stryker, 2017). The distinct effects that sound and locomotion have on visual responses also adds nuance to our understanding of how motion affects visual processing, as other groups have predominantly used responses averaged across the duration of the stimulus presentation in categorizing motion responsive neurons in V1 (Neil and Stryker, 2010; Dadarlat and Stryker, 2017). Our findings indicate that the timing of cross-sensory interactions is an important factor in the classification and quantification of multisensory effects.

We also observed that motion decreases the magnitude of the enhancing effect that sound has on the onset of the visual response (Figure 6G,H, Figure 7F,H). This finding suggests a degree of suppressive effect that motion has on this audiovisual interaction. A potential mechanism for this result may relate to the circuits underlying audiovisual integration in V1. Other groups have shown using retrograde tracing, optogenetics and pharmacology that the AC projects directly to V1 and is responsible for the auditory signal in this region (Falchier et al., 2002; Ibrahim et al., 2016; Deneux et al., 2019). It is currently understood that unlike in V1, in other primary sensory cortical areas including the A1, movement suppresses sensory evoked activity (Nelson et al., 2013; Schneider and Mooney, 2018; Bigelow et al., 2019). Therefore, one explanation for this observation is that despite motion enhancing the visual response magnitude in the absence of sound, the suppressive effect that motion has on sound-evoked responses in the AC leads to weaker AC enhancement of visual activity on trials in which mice display robust movement. A detailed experimental approach using optogenetics or pharmacology would be required to test this hypothesis of a tripartite interaction and would also reveal the potential contribution of other auditory regions.

### Enhanced response magnitude and consistency combine to improve neuronal encoding

Signal detection theory indicates that improved encoding can be mediated both by enhanced signal magnitude as well as reduced levels of noise (von Trapp et al., 2016). When using purely magnitude-based metrics of discriminability, OSI and DSI, we found a small reduction from the visual to audiovisual conditions (Figure 4E,F). However, we also observed that sound reduced the CV of visual responses (Figure 5), a measure of the trial-to-trial variability in response. When we measured the *d’* sensitivity index of neuronal responses, a measure that factors in both the mean response magnitude and trial-to-trial variability, we found that sound improved the discriminability of drifting grating orientation and direction (Figure 8A,B). These findings indicate that the improved discriminability of visual responses in individual neurons was mediated not only by changes in response magnitude but also by the associated improvement in response consistency between trials, despite the mild sound-associated reduction in OSI and DSI. The relevance of trial-wise variability is further supported by our observation of reliably orientation- and direction-selective neurons despite relatively low OSI and DSI (Figure 4A-D), a metric agnostic to response variability. Prior studies using patch-clamp primarily in anesthetized animals approaches showed that V1 neurons sharpen their tuning profiles in response to sound a magnitude-based coding scheme, with some degree of recapitulation in the awake state (Ibrahim et al., 2016). The difference between these findings and those reported in the current study potentially indicate different coding schemes present in anesthetized and awake brains, additionally modified by unrestricted uninstructed movement during both visual and audiovisual trials, of which both factors are known to affect cortex-wide neuronal dynamics (Musall et al., 2019). It is therefore important to consider response variability in awake brains in addition to magnitude-based metrics when quantifying tuning and discriminability in neurons (Churchland et al., 2011; Mazurek et al., 2014).

Multiple studies found variable effects of sound presentation on visual responses in V1 (Meijer et al., 2017; McClure and Polack, 2019). Dependent on selection criteria of stimulus responsiveness, stimulus parameters, and visual contrast level, both studies found that sound presentation evoked either enhancement or suppression of V1 activity. Importantly, these studies found that highly selective neuronal responses to visual stimuli are enhanced by sounds at intermediate visual contrast, which is in agreement with our analysis. Some differences could potentially be attributed to differences in sound presentation and stimulus selection. Meijer et al., 2017 finds that visual responses are enhanced on average by sound stimuli that are congruent with the visual stimulus. It is possible that the congruent stimulus for the majority of neurons approximated the neurons’ preferred temporally frequency, which activated neurons similarly to the white noise burst in our study. Meijer et al., 2017 also used a white noise burst and found that this auditory stimulus enhanced neuronal responses at intermediate visual contrast levels and suppressed them at high visual contrast levels. The difference between our results and those by McClure and Polack, 2019 may also be attributed to differences in the sound stimulus: they used pure tones that would activate neurons tuned to that particular frequency. Conversely, the duration of our white noise was matched to the visual stimulus, which were also longer in our study, therefor recruiting neurons in V1 across layers. Combined, our findings build on these two studies to explicitly test the role of locomotion and its sound-evoked qualities, providing a deeper understanding of the effects of locomotion through an expanded generalized linear model and of changes in variability as the key component to improved population-level decoding.

Inhibitory and disinhibitory effects may arise at different points within V1 with distinct effects on neuronal dynamics. Iurilli et al., 2012 found that presentation of sound burst drove a suppressive effect in V1, driven by cortico-cortical excitatory projections from AC to infragranular neurons. The effects in supra-granular layers were predominantly suppressive. We note that this study differed from ours in large part because of the visual stimulus. Iurilli et al., 2012 presented a brief light flash, which would activate V1 neurons in a different fashion than the stimulus that ours and later studies used, a prolonged drifting grating. Whereas a single flash may evoke wide-spread adaptation and suppression in V1, a drifting grating is a stimulus that targets specific V1 neurons depending on their orientation. Furthermore, we observed that sound may suppress the sustained portion of the visual response in the absence of motion (Figure 7F), suggesting that sound-induced excitation and inhibition may be temporally dependent as well.

### Stimulus parameters relevant to audiovisual integration

Sensory neurons are often tuned to specific features of unisensory auditory and visual stimuli, and these features are relevant to cross-sensory integration of the signals. In the current study we paired the visual drifting gratings with a static burst of auditory white noise as a basic well-controlled stimulus. Indeed, it is known that the audiovisual stimulus profile affects the degree to which sound is integrated with the visual signal (Bizley et al., 2016; Meijer et al., 2017; Atilgan et al., 2018). Previous studies found that temporally congruent audiovisual stimuli, e.g. amplitude-modulated sounds accompanying visual drifting gratings, evoke larger changes in response than temporally incongruent stimuli in the mouse visual cortex (Meijer et al., 2017; Atilgan et al., 2018), and therefore using such stimuli would potentially result in even stronger effects than we observed. However, in other brain regions such as the inferior colliculus, audiovisual integration is highly dependent on spatial congruency between the unimodal inputs (Bergan and Knudsen, 2009). Additional studies are needed to explore the full range of auditory stimulus parameters relevant to visual responses in V1.

Our results show that spatially congruent, static white noise is sufficient to improve V1 neuronal response magnitude and latency to light-evoked responses. These results likely extend to natural and ethologically relevant stimuli as well. Indeed, rhesus macaque monkeys demonstrate psychometric and neurometric improvements in tasks such as conspecific vocalization detection and object recall (Hwang and Romanski, 2015; Bigelow and Poremba, 2016; Breman et al., 2017). Humans are also capable of perceptually integrating audiovisual stimuli ranging from paired visual drifting gratings and auditory white noise (Lippert et al., 2007; Chen et al., 2007), to the McGurk effect and virtual reality simulated driving (McGurk and MacDonald, 1976; Marucci et al., 2021). We therefore posit that the audiovisual integration of basic sensory stimuli in early sensory areas may form the foundation for functional integration by higher cortical areas and ultimately behavioral improvements.

### Neuronal correlates of multisensory behavior

Our findings of multisensory improvements in neuronal performance are supported by numerous published behavioral studies in humans and various model organisms (Gleiss and Kayser, 2012; Meijer et al., 2018; Stein et al., 2020). Training mice to detect or discriminate audiovisual stimuli allows the generation of psychometric performance curves in the presence and absence of sound. We would hypothesize that the intermediate visual contrast levels in which we see improvements in neural encoding would align with behavioral detection threshold levels. One could also correlate the trial-by-trial neural decoding of the visual stimulus with the behavioral response on a stimulus discriminability task, an analysis that could provide information about the proximity of the V1 responses to the behavioral perception and decision. Additionally, a behavioral task could allow the comparison of neural responses between passive and active observing, helping to reveal the role of attention on how informative or distracting one stimulus is about the other.

### Multisensory integration in other systems

It is useful to contextualize audiovisual integration by considering multisensory integration that occurs in other primary sensory cortical areas. The auditory cortex contains visually responsive neurons and is capable of binding temporally congruent auditory and visual stimulus features in order to improve deviance detection within the auditory stimulus (Atilgan et al., 2018; Morrill and Hasenstaub, 2018). Additionally, in female mice, pup odors reshape AC neuronal responses to various auditory stimuli and drive pup retrieval behavior (Cohen et al., 2011; Marlin et al., 2015), demonstrating integration of auditory and olfactory signals. However, whether these forms of multisensory integration rest on similar coding principles of improved SNR observed in the current V1 study is unknown. Investigation into this relationship between the sensory cortical areas will help clarify the neuronal codes that support multisensory integration, and the similarities and differences across sensory domains.

## Acknowledgements

The authors thank Gabrielle Samulewicz for assistance with experiments and members of the Geffen laboratory for helpful discussions, as well as Dr. Jay Gottfried and Dr. Yale Cohen at the University of Pennsylvania. This work was supported by funding by the National Institute on Deafness and Other Communication Disorders at the National Institute of Health grants 5T32DC016903 to AMW, F31DC016524 to CFA, and R01DC015527, R01DC014479, and R01NS113241 to MNG.

